# The nutritional value of invertebrate aquatic foods

**DOI:** 10.1101/2025.03.26.645521

**Authors:** Jessica Zamborain-Mason, Nisha Marwaha, Seo-Hun Yoo, Christina C. Hicks, James P.W. Robinson, Luisa R. Abucay, Laura G. Elsler, Jacob G. Eurich, Whitney R. Friedman, Jessica A. Gephart, M. Aaron MacNeil, Julia G. Mason, M.L. Deng Palomares, Vina A. Parducho, Katherine L. Seto, Kristin M. Kleisner, Daniel F. Viana, Christopher D. Golden

## Abstract

Aquatic invertebrates are a diverse, nutrient-dense, and socio-ecologically important food whose contribution to human nutrition is frequently overlooked. We quantify their contribution to global nutrient supplies and estimate the nutrient content of >50,000 macroinvertebrate species. Current aquatic invertebrate production supplies the equivalent annual requirement for >6 billion people in terms of vitamin B12 and selenium; >1 billion people for copper, omega 3 fatty acids, iodine and zinc; and >100 million people for nutrients such as vitamins B2 and B3, iron, manganese, and magnesium. Nutrient composition differs among taxonomic groups, consumption patterns, and environmental and life-history factors. Our study highlights the benefits of integrating aquatic invertebrates into dietary portfolios across global societies, mainstreaming their nutritional importance in development projects, sustainability assessments and food policy.

## Main Text

The majority of the global population has inadequate micronutrient intake (*1*), leading to cascading adverse effects on economies and human health (*2*). Meanwhile, increasing anthropogenic pressures are expected to drive micronutrient declines in major food sources (*3, 4*), highlighting the need to better understand and account for the nutrient delivery from typically under-appreciated foods. Invertebrates account for most of the animal biomass and diversity on earth (*5*). Aquatic invertebrates, more specifically, are diverse (>1 million estimated species; (*6*)), culturally and socially important (*7*), ecologically significant (*8*), economically valuable (e.g., invertebrate ex-vessel prices can reach >10,000 $USD/ton; (*9*)), and highly nutritious (*10*). For example, aquatic invertebrates such as molluscs have higher nutrient concentrations per 100g than other aquatic foods such as ray-finned fishes (e.g., vitamin B_12_, iron, iodine, manganese, magnesium, or zinc; (*11*)). Furthermore, access to their diversity can help buffer current and future diet-related non-communicable diseases and nutritional deficiencies (*10,12*), especially in the context of environmental change (*4,13*).

Yet, despite their established importance, most aquatic invertebrates are grossly underrepresented in fishery (*14,15*), conservation (*16–18*), and nutritional assessments (e.g., *19-22*). For example, key advancements in ray-finned fish research have allowed the discernment of ecological, environmental (*20*), and phylogenetic traits (*19*) associated with nutrient content, making the prediction of nutrient composition for finfishes lacking such data possible (*20*). Among others, these advancements have allowed estimating the potential contribution of fish-based food strategies to global nutrition-security (*20*). Yet, similar research for invertebrates has lagged, despite their diversity (*6*), their increasing production levels relative to finfish (*14*), and their strong potential as a productive and low environmental footprint food (*23,24*).

Understanding the contribution of aquatic invertebrates to nutrient supplies, and their ability to meet nutritional adequacy targets, is crucial to (i) provide empirical evidence of current and potential nutritional contributions of aquatic invertebrates, and (ii) understand the nutritional and public health implications of effective invertebrate monitoring, assessment, management and policy to meet multiple sustainability targets (e.g., Sustainable Development Goal of “zero hunger”, “life below water” or “gender equality”).

Here, we integrated the Aquatic Food Composition Database (*10*) with marine capture fisheries (*25*), aquaculture production (*26*) and species-level environmental and ecological trait data (*27*) to estimate the contribution of aquatic invertebrates to global nutrient supplies (i.e., available pool of nutrients produced or captured). First, we quantified the relative contribution of invertebrates to nutrient supplies of global marine capture fisheries and aquaculture production, identifying the sectors, species groups, and countries that provide most aquatic invertebrate nutrients to human populations. In our analyses, we estimate global fisheries and aquaculture production of 29 nutrients essential for human health (Methods; *11*). Second, we developed a series of Bayesian hierarchical models to determine the variability and potential drivers of nutrient content in aquatic invertebrates. For this, we combined species-level environmental and ecological trait data (*27*) with an updated invertebrate species-specific nutrient composition database (e.g., *10*) that includes 16,481 samples from 471 invertebrate species. Finally, we leveraged model predictions to estimate the nutrient composition of 50,807 macroinvertebrate species registered globally (*27*), showcasing their potential contributions to public health and regional food security.

### Nutrient contributions of aquatic invertebrates

First, combining global statistics on marine capture fisheries and aquaculture production with the nutrient content of aquatic foods, we estimated the contribution of aquatic invertebrates to global aquatic animal source nutrient supplies. We found that several nutrients were more concentrated in aquatic invertebrates than in other taxonomic groups, such as finfish (e.g., zinc; (*11*)). As a result, invertebrates contributed disproportionately to supply of key essential nutrients, relative to their production volume. For marine capture fisheries, invertebrate aquatic foods contributed relatively more nutrient supply than biomass supply for 11 of the 29 nutrients examined (Fig. 1A). More specifically, while invertebrates contributed ∼16% of total marine capture volumes, they comprised: >40% of copper and zinc nutrient supplies from marine catches; >25% of manganese, vitamin B_9_ (folate), sodium and vitamin C; and >16% of selenium, magnesium, vitamin E, phosphorus and iron supplies (i.e., relatively more nutrient supply than biomass supply).

**Fig. 1.**
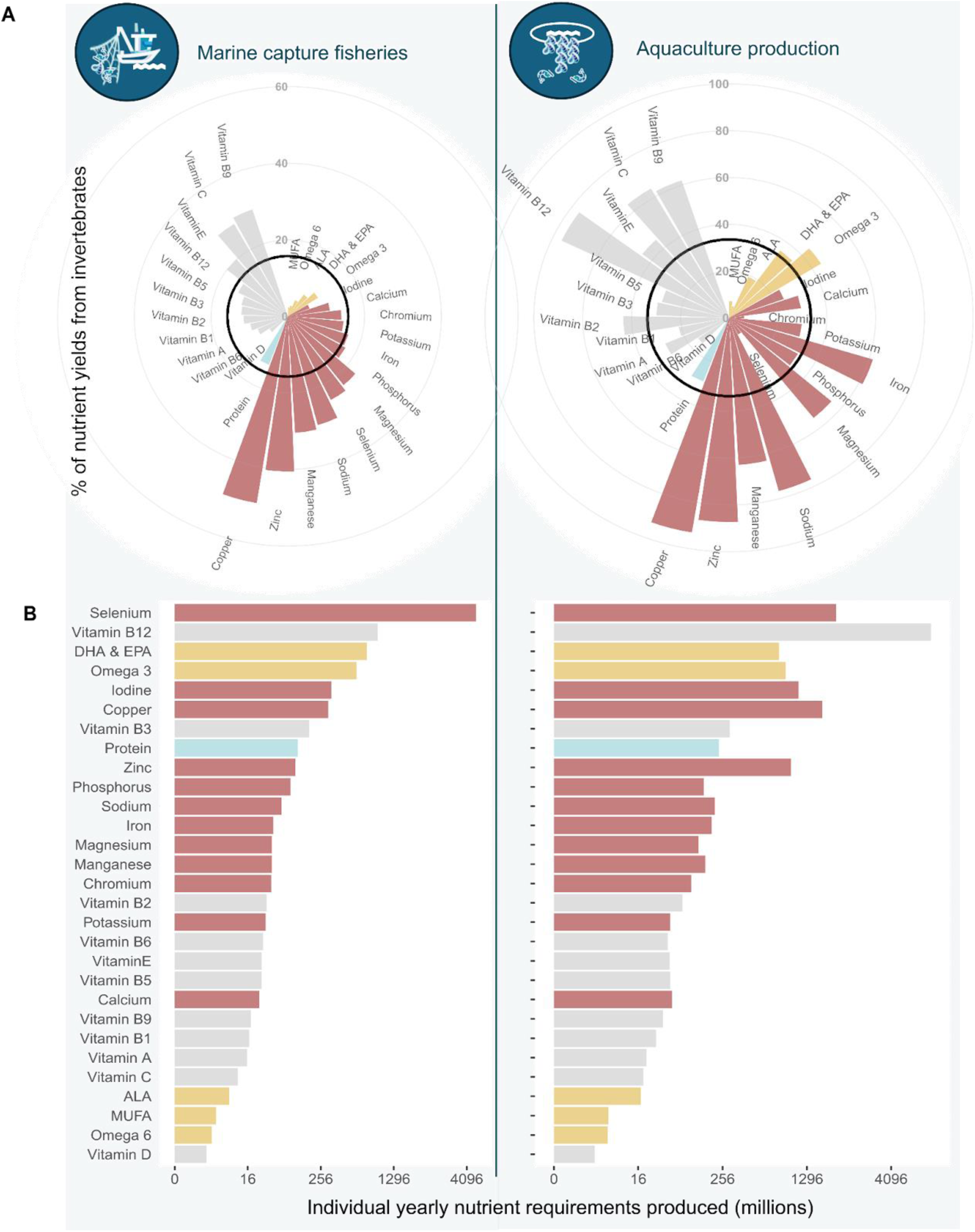
Nutrient contribution of aquatic invertebrates. **(A)** Percent of nutrient yields coming from invertebrates for marine capture fisheries (left) and aquaculture production (right). Black circles indicate the percent contribution of invertebrates in terms of marine capture or aquaculture production volumes (i.e., live weight). Bars surpassing the black line means that invertebrates contributed disproportionately more nutrients relative to catch or production volumes. See Fig. S1 for the figure using edible weights. **(B)** Number (in millions) of yearly nutritional requirements produced from aquatic invertebrates in marine capture fisheries or aquaculture production for 2019. For each nutrient, we use the average requirements across demographic groups (Table S3). To aid clarity the x axis was fourth-root transformed and reported values are on an arithmetic scale. Bars in **A-B** are color-coded by nutrient type: minerals in red, fatty acids in yellow, protein in blue, and vitamins in grey. See Figs. S2-S5 for country, sector and phylum specific variability in invertebrate contributions.

Within aquaculture production, relative to production volumes, invertebrates disproportionately contributed to the supplies of 13 nutrients (Fig 1A). While invertebrates contributed ∼ 34% of total aquaculture production volumes in 2019, they contributed: >70% of aquaculture copper, zinc, sodium and vitamin B_12_ (cobalamin); >60% of manganese, iron, vitamin B_9_ and vitamin C; and >35% of magnesium, omega 3 fatty acids including docosahexaenoic acid and eicosapentaenoic acid (DHA & EPA), vitamin B_2_ (riboflavin) and vitamin E. Note that, due to the species composition differences, aquacultured invertebrates contributed more nutrients (relative to production volumes) than marine capture invertebrates, with a different ranking and composition of nutrient contribution (Fig.1A). In both marine capture fisheries and aquaculture production, we found that the relative importance of nutrient supply was robust to whether we accounted for live weight or edible weight (Fig. S1; Methods). Other nutrients, such as potassium, chromium, calcium, and iodine, vitamins B_1_ (thiamin), B_3_ (niacin), B_5_ (pantothenic acid), B_6_, A and D, protein and omega 6, and alpha-linolenic acid (ALA) fatty acids, were primarily supplied by non-invertebrate aquatic foods (i.e., invertebrates contributed less to these nutrient supplies when compared to their contributions in terms of volumes; Fig.1A). This emphasizes the role that a diverse portfolio of aquatic foods can have in tackling nutrition insecurity more broadly (*10*).

Second, to determine the public health implications of aquatic invertebrate nutrient supplies, we calculated the number of yearly recommended nutrient intakes (RNI; recommended dietary allowance (RDA) or adequate intake (AI); Methods) potentially met from aquatic invertebrate production. We found that from the list of 14 nutrients highlighted in Fig. 1A, invertebrates met the highest number of recommended nutrient intakes for vitamin B_12_, selenium, copper, omega 3 fatty acids including DHA & EPA, zinc, sodium, phosphorus, iron, manganese, magnesium, vitamin B_2_, vitamin E and vitamin B_9_ (in order of overall importance; Fig. 1B). More specifically, we found that invertebrate production from aquaculture and marine capture fisheries in 2019 supplied the equivalent annual requirements of >6 billion people for vitamin B_12_ and selenium; >1 billion people for copper, omega 3 fatty acids including DHA & EPA, iodine and zinc; >300 million people for vitamin B_3_ and protein; >100 million people for sodium, phosphorous, iron, manganese, magnesium, chromium and vitamin B_2_; >50 million people for potassium, calcium, vitamins B_5_ E, B_6_, B_9_ and B_1_; >20 million people for vitamins A and C, and ALA; and >1 million people for monounsaturated fatty acids, omega 6 fatty acids and vitamin D (Fig. 1B). Dietary contributions varied by production method, with marine capture fisheries producing more selenium (4.6 billion yearly requirements) and DHA & EPA (765.9 million), and aquaculture producing more vitamin B_12_ (6.4 billion), copper (1.6 billion), iodine (1.1 billion) and zinc (1 billion). Overall, our study demonstrates that aquatic invertebrates are likely providing substantial health benefits to society, provided their nutrient supplies are sustainable, well distributed and accessible to those who most need them.

Third, disaggregating capture and production by sector (i.e., industrial, artisanal, subsistence, or recreational fishing for marine capture fisheries; marine or freshwater for aquaculture production), taxa and location (e.g., country) revealed high variability in invertebrate contributions to nutrient supplies. In marine capture fisheries, the industrial sector supplied most of aquatic invertebrates in terms of total capture volumes (64%), followed by the artisanal and subsistence sectors (32 and 3%, respectively). Due to the high volumes captured, the industrial sector produced the highest amount of nutritional requirements (with the exception of selenium; Fig. S2). However, relative to sector-specific catch volumes (e.g., including fish), invertebrate catches and nutrients had the highest contributions to artisanal and subsistence sectors (Fig. S3). For example, subsistence invertebrate fisheries contributed disproportionately more nutrients than catch volume (i.e., 16 of the 29 nutrients; Fig. S3), and artisanal invertebrate fisheries provided the highest yearly requirements of selenium in 2019 (>3 billion yearly requirements; Fig. S2). From the six phyla recorded (i.e., mollusca, arthropoda, echinodermata, cnidaria, porifera and annelida), molluscs and arthropods dominated invertebrate catches and nutrient supplies, with captured molluscs supplying (in 2019) the equivalent annual requirements for > 4 billion people for selenium and > 400 million people for vitamin B_12_ and DHA & EPA (Fig. S2), and disproportionately contributing to 21 of the 29 nutrients, relative to their phyla-specific catch volume (Fig. S4).

Invertebrate contributions in marine capture fisheries also differed by country. Exclusive economic zones (EEZ) of China, Japan and Vietnam caught the highest invertebrate volumes, with China alone producing 5 and 12 % of the total yearly requirements of selenium and vitamin B_12_ caught globally (248 and 137 million, respectively). Other exclusive economic zones with lower total invertebrate catch, such the Caribbean EEZ of Honduras (i.e., 0.38% of China’s catch), had higher prevalence of invertebrates in their catches (>85%), for instance capturing the equivalent annual selenium requirements for >53 million people. Several tropical island developing states were notable in having high invertebrate contributions to nutrient supply, with Fiji, Kiribati and Tonga providing more nutrients than expected from catch volume (for 22 out of 29 nutrients; Fig. S5)

In aquaculture production, most of the invertebrates were supplied by the marine sector (87% in terms of live weight). Invertebrate mariculture in 2019 provided 76% of total mariculture supply, and > 85% of marine cultured zinc, copper, manganese, sodium, vitamin B_12_, magnesium, folate and iron supplies (Fig. S3). This sector alone produced > 4 billion yearly requirements of vitamin B_12_ (Fig. S2). Inland invertebrate aquaculture accounted for a small portion of inland aquaculture (7.2 %) but was nutrient rich, disproportionately contributing to 19 of 29 nutrients (i.e., higher nutrient yields than expected from inland production volumes; Fig. S3). Similar to marine capture fisheries, cultured molluscs and arthropods also contributed most invertebrate nutrients, with cultured molluscs (i) dominating nutrient supplies when compared to production volumes (21 out of the 29 nutrients examined; Fig. S4) and (ii) supplying the equivalent annual requirements for > 6 billion people for vitamin B_12_ and >1 billion people for selenium, copper and iodine (Fig. S2).

China, Vietnam and Indonesia produced most invertebrate aquaculture, with China producing 88%, 74% and 82% of the total yearly requirements of vitamin B_12_, selenium and copper cultured from invertebrates globally (i.e., 5.6, 1.5 and 1.3 billion yearly requirements, respectively). In other countries such as Ecuador, despite relatively low production volumes in comparison to China, invertebrates accounted for >95% of cultivated aquatic foods, producing >10 million requirements of selenium, vitamin B_12_ and B_3_. Relative to country-specific production volumes, the republic of Moldova ranked as the country with greater nutrient contributions from cultured invertebrates, with 21 nutrients out of 29 being disproportionately contributed by invertebrates in comparison to other aquatic foods (Fig. S5). Combined, all these sector, phyla and country-specific results highlight that nutrient-sensitive approaches to invertebrate and aquatic food management in different geographies need to consider specific fishery and aquaculture contexts (e.g., account for the variability in species composition profiles and quantities being caught or produced).

### Nutrient content variability in aquatic invertebrates

Given the nutritional importance of invertebrates (e.g., Fig. 1), and building on predictive frameworks developed for finfishes (*20*), we estimated the variability and potential drivers of invertebrate nutrient content using species-specific invertebrate nutrient composition data, ecological and environmental trait data, and a series of Bayesian hierarchical models (Methods; Table S1). First, we found that some nutrients vary considerably more than others among samples and taxonomic groupings (Fig. S6). For example, variance decomposition results from our Bayesian hierarchical models show that selenium (for minerals) and vitamins B_12_ and A (for vitamins) had the highest variability. Similarly, for macronutrients (i.e., fatty acids and protein), alpha-linolenic acids (ALA) within invertebrates varied substantially more than others such as protein or omega 6 fatty acids. These variability differences highlight the nutrients that may be best prioritized in future studies, for example, to model and quantify spatio-temporal variability in the nutrient content of aquatic foods.

Second, we show that, when controlling for environmental and ecological traits, some taxonomic groups have higher nutrient concentration than others, but not consistently across all nutrients (Fig. 2A, Fig. S7). For example, compared to average invertebrate values (Fig. 2A), bivalves had the highest concentrations of iron and vitamins B_9_, B_2_ and B_12_, whereas other groups such as echinoderms (e.g., sea urchins and sea cucumbers) had the highest concentrations of magnesium, manganese and sodium (Fig. 2). These group-specific differences in nutrient content could help inform public health programmes and policies aimed at harnessing invertebrate foods to improve undernourishment, particularly in locations where policies may target specific nutrient deficiencies and vulnerable populations.

**Fig. 2.**
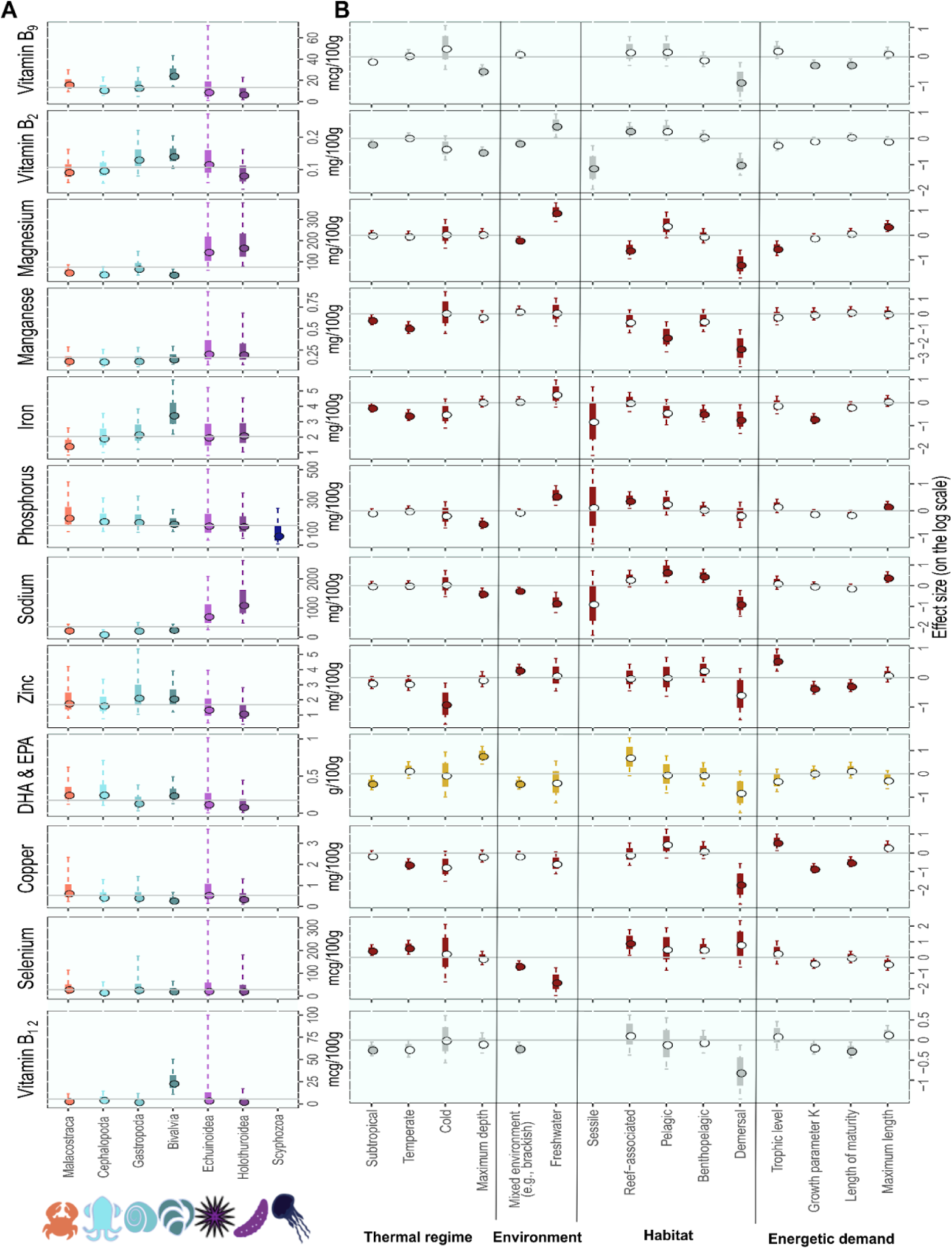
Variability and potential drivers of nutrient content in aquatic invertebrates. **(A)** Estimated nutrient concentration per taxonomic class. Points are the median estimated nutrient concentration for that class (i.e., global intercept plus relevant phylum and class intercept offsets), solid and dashed lines represent 50 and 90% uncertainty intervals, respectively. Vertical line represents the median estimated global intercept for each nutrient across all invertebrates. Symbols are color-coded by class group. **(B)** Effect sizes of different ecological and environmental factors (Table S1) on average nutrient concentrations (in log scale). Points are medians, and the solid and dashed lines represent 50 and 90 % uncertainty intervals, respectively. Horizontal lines are set at 0 (i.e., the baseline category). Nutrients are color-coded by nutrient type: minerals in red, fatty acids in yellow, protein in blue, and vitamins in grey. If the parameter 90% uncertainty intervals overlapped zero, the point is not filled. Estimates in both **(A-B)** are baselined for raw muscle tissue at average environmental and ecological continuous conditions (e.g., depth, maximum length, length of maturity, growth parameter K and trophic level) and most common categories (e.g., tropical, marine, and benthic). Missing estimates (e.g., Sciphozoa in **A** or sessile in **B**) indicate a lack of species-specific nutrient composition data under that category. Note that while we developed models for all nutrients with sufficient species-specific invertebrate data (i.e., all but vitamin D and E; Methods), this figure highlights nutrients that were simultaneously (i) disproportionately contributed by invertebrates, and (ii) contributed the most to meeting human nutritional adequacy (e.g., Fig 1). See Fig. S7 for the remaining nutrients and Fig. S8 for nuisance parameter results (i.e., how nutrient concentrations vary with food part or processing sampled).

Third, similar to finfish (*20*), we found that several ecological and environmental predictors had strong associations with nutrient concentrations in aquatic invertebrates (Fig. 2B; Fig. S7). Demersal invertebrate species, which live and feed in the water column near the seafloor, for example, consistently exhibited lower nutrient concentrations compared to reef-associated species, which had relatively higher concentrations across key nutrients examined (e.g., for iron, zinc, sodium, copper, manganese and vitamins B_2_, B_9_ and B_12_, >2 times the effect size; Fig. 2B). Similarly, invertebrate species sampled from freshwater environments showed higher vitamin B_2_, magnesium, and phosphorus concentrations and lower sodium and selenium concentrations compared to species sampled from marine or mixed (e.g., brackish) environments (e.g., median and 90% uncertainty effect sizes did not overlap zero; Fig. 2B). We also found nutrient concentration patterns with the latitudinal environment (e.g., thermal regime) occupied by the species, with cold and tropical species sometimes having similar nutrient concentrations but subtropical and temperate aquatic invertebrates showing either relatively higher (e.g. selenium) or lower (e.g., manganese, iron, and vitamin B_12_) nutrient concentrations per 100g of raw muscle tissue (Fig. 2B). Species that inhabit deeper waters were also richer in DHA & EPA but had lower concentrations of vitamins B_9_, B_2_, phosphorus, and sodium than species that reach lower maximum depths. These trends, besides giving some potential indication of the nutrient sources (e.g., freshwater streams typically have higher anthropogenic minerals that could be absorbed by organisms (*28*)), suggest that the accumulation of nutrients in the muscle tissue of invertebrates is likely influenced by the thermal regime the species is exposed to, with some nutrients potentially having temperature-specific optima (*29*).

Fourth, we show that different scenarios of consumption patterns, for example, food processing methods and body parts potentially consumed, can shape the nutrient concentration of aquatic foods. Part of our approach aimed to control for factors (e.g., food processing and body parts) in our data that could impact the nutrient composition of aquatic invertebrates ((*30*); Methods; Table S1). We found these factors had strong associations with the nutrient content in aquatic invertebrates, often of higher magnitude than ecological and environmental traits (Fig. S8). For example, whole and dried invertebrates tended to have higher nutrient concentrations (e.g., calcium, zinc, sodium, potassium, phosphorus, or magnesium) than raw muscle tissue (Fig. S8). These results highlight the need to consider what (i.e., edible portion; (*31*)) and how (i.e., processing and preparation) aquatic foods are eaten in different dietary cultural contexts (*32*). Further, these results could indicate opportunities for value-added processing where highly nutritious body parts are not generally consumed (e.g., “fish crisps” from crab shell waste; (*33*)).

### Nutrient composition estimates for data-limited invertebrates

Most aquatic foods produced by aquaculture and marine capture fisheries lack empirical species-specific nutrient composition data (66-97% dependent on nutrient; Fig. S9). Thus, to advance research on the nutrient potential of aquatic invertebrates, we used model outputs to provide nutrient composition estimates for a total of 50,807 potentially edible macroinvertebrate species registered globally with available taxonomic, environmental and/or life-history traits (i.e., SeaLifeBase (*27*); Methods). This revealed four key findings. First, based on mean values across all aquatic macroinvertebrate species, 100 grams of raw muscle tissue can provide > 95% of selenium, chromium and vitamin B_12_ daily requirements, 67% of copper, 48% of niacin, 38% of zinc and >25% of protein, magnesium, and iron daily requirements (Fig. 3A). Second, for some nutrients (e.g., copper, zinc, or magnesium), our models predicted high variation among invertebrate species in the human requirements provided per 100g of muscle tissue, whereas for others (e.g., vitamin B_6_ or B_9_) there was less species-to-species estimated variability (Fig. 3A). Third, by focusing only on species which have a defined fisheries importance category (Methods; Fig. S10), we showed that, based on mean values, invertebrate species typically used for subsistence are estimated to be relatively richer in iron, whereas highly commercial species are relatively richer in copper and zinc, and bycatch species are relatively richer in vitamin B_3_ (Fig. S10).

**Fig. 3.**
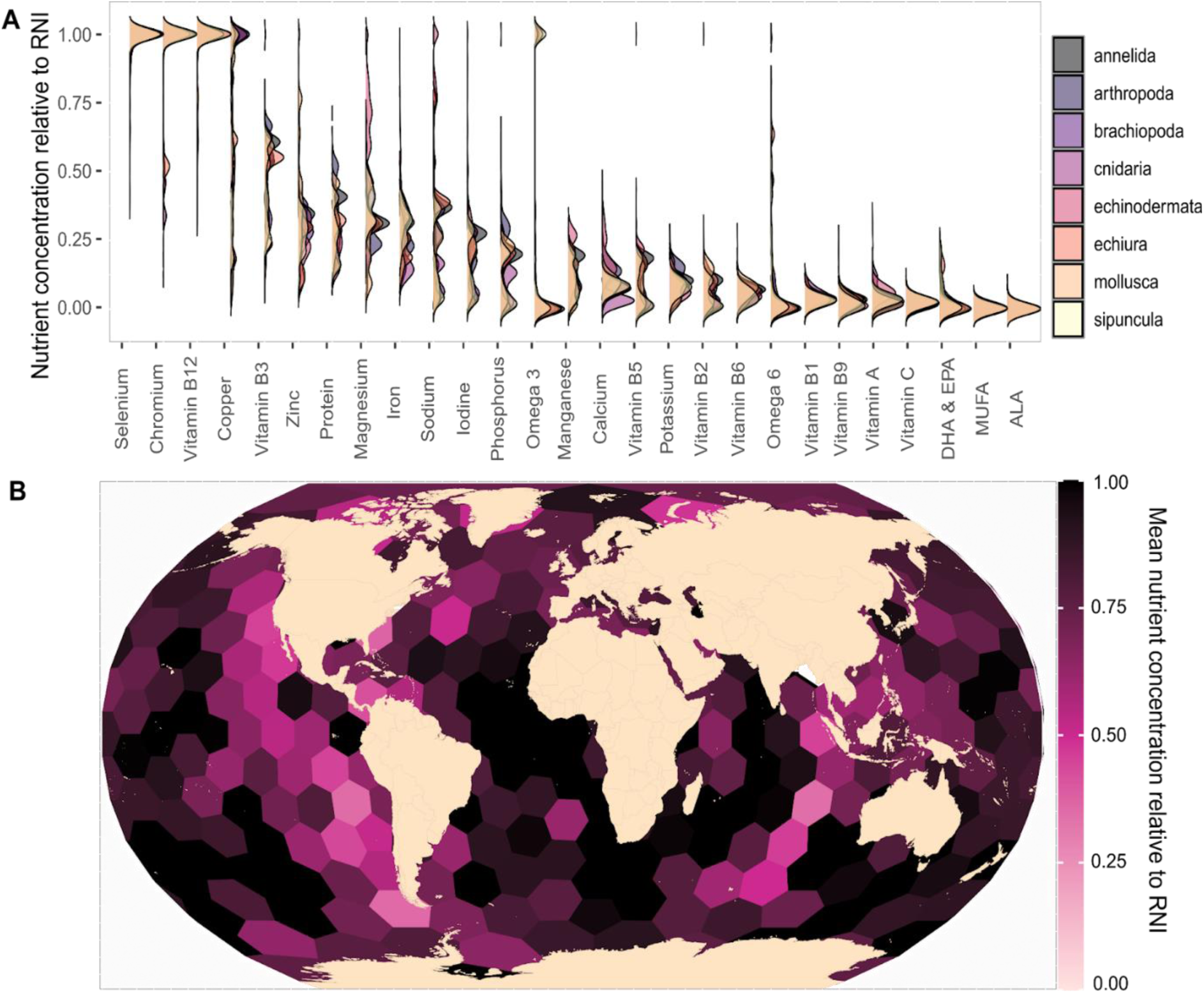
Estimated nutrient concentrations relative to recommended nutritional intakes (RNI) for potentially edible macroinvertebrates. **(A)** Median nutrient concentration predictions per 100g of raw muscle tissue relative to recommended dietary intakes for a total of 50,807 potentially edible macroinvertebrate species available in SeaLifeBase (*27*). Densities are color-coded by phylum groups. **(B)** Geographic distribution of potentially edible aquatic macroinvertebrates based on species occurrences. Hexagons are bins of equal area that represent the mean nutrient concentration per 100g of muscle tissue relative to daily RNI (across nutrients). See Fig. S11 for individual nutrients. If values in **A-B** were above one we restricted them to one.

Finally, by matching estimated nutritional values to species occurrence data ((*34*); Methods), we show the geographic distribution of nutrient content in aquatic invertebrates (e.g., nutrient requirements per 100g of muscle tissue, averaged across all nutrients which we developed models for; Fig. 3B). This revealed that available aquatic invertebrates have great potential for many regions with high inadequate intake of nutrients (*1*). For example, the prevalence of inadequate zinc intake is high in South Asia, the Middle East and Africa (>75% of the population; (*1*)) but the invertebrate species that occur in such regions have high zinc concentrations (Fig. S11), with 100g of invertebrate muscle tissue providing >50% of daily zinc intake requirements (i.e., 8.91 mg which is the average daily zinc requirements across demographic groups; Methods). Overall, these model-based aquatic invertebrate nutrient predictions can inform current and future developmental projects, aquatic food management and policy, and also increase our understanding on the potential trade-offs that may exist between nutrient supplies and the risk of toxins and contaminants (*35*).

### Aquatic invertebrates to meet global nutrition and sustainability targets

Our study quantified the contribution of aquatic invertebrates to human nutrition and estimated the variability and potential drivers of invertebrate nutrient content. In the process, we developed predictive models to estimate nutrient concentrations of aquatic invertebrates when analytical data are not available. This section highlights three future research avenues that can build upon our work to further advance aquatic invertebrate accounting, facilitating the acceleration of theory and practice to improve nutrition security and achieve multiple global targets, such as the Sustainable Development Goals.

The first research avenue is the need for better empirical aquatic invertebrate nutrient composition data. Models that predict the nutrient composition of aquatic foods using phylogenetic, taxonomic, and environmental and ecological trait information (e.g., (*19,20*); Fig. 2 in the present study) are useful tools to fill in data availability gaps, develop understanding of how environmental changes are likely to affect the nutrients available (*4*), and help to steer aquatic food systems towards greater sustainability (*22*), equity (*36*), and nutrition security (*20*). For example, available predictions from our work (e.g., Fig. 3) can be used to inform targeted location-based research to aid nutrition interventions (e.g., incorporating invertebrates in diets to address specific nutritional adequacies; (*37*)), nutrition-sensitive approaches to fisheries management (*22*), or research exploring the sustainability and climate-resilience of aquatic food systems (*36*). However, models, while useful, are only as good as the data that informs them. For example, our study analyzed species-specific nutrient composition data for 471 invertebrate species, yet the number of species varied by nutrient (Table S2). As a consequence, some edible macroinvertebrate groups such as jellyfish (e.g., Cnidaria) were absent from our estimates of invertebrate contributions to aquatic nutrient supplies (Fig. 1), and scarce (i.e., low sample size, mainly informed by estimated intercepts and ecological trait patterns of other groups) in our global models of nutrient concentration (e.g., Figs. 2-3). Similarly, some nutrients that are essential in diets and rich sources in aquatic foods, either had limited sample size (e.g., total omega 3 fatty acids; Table S2) or lacked sufficient species-specific invertebrate nutrient composition data to perform model predictions (e.g., vitamins E and D; Table S2). Future work that samples aquatic invertebrates for their nutrient content (and consistently reports it; e.g., relative weight; Table S1) could target specific nutrients (e.g., vitamins D and E), specific aquatic invertebrate groups (e.g., edible sea worms or jellyfish), and invertebrates with specific ecological and environmental trait data (e.g., freshwater molluscs) to increase aquatic invertebrate representation in model predictions, eliminate potential data structures in currently available data (e.g., allow for covariate balance in predictive tools; (*38*)) and improve overall understanding of aquatic invertebrate nutrients.

The second, and related, research avenue is that which accounts for intra- and inter-specific spatio-temporal variability in aquatic invertebrates, and aquatic foods more broadly. We used an updated version of the Aquatic Food Composition Database (*10*) to assign nutrient composition estimates to individual species and matched this to species-specific ecological and environmental traits in SeaLifeBase (*27*). While these are the largest empirical compilations available of aquatic food nutrient composition and ecological information, spatio-temporal nutrient composition data (e.g., how nutrients vary spatially and temporally within species, or temporally across species) was limited (Table S1). However, our results show which nutrients vary most (e.g., selenium or vitamin B_12_; Fig. S6) and which nutrients show patterns with spatio-temporal variables (e.g., selenium residuals and year of sampling; Fig. S16), and thus could be used to prioritize research that evaluates the drivers of nutrient content in aquatic foods. For example, the environmental drivers of nutrient content (e.g., how nutrient composition is likely to vary with temperature and ongoing climate change; (*39*)), or how ontogenetic shifts (e.g., changes in habitat; (*40*)) or biogeochemical cycles (*28*) influence the nutrient pool available from aquatic foods under different contexts.

Finally, given the food and nutrition-security relevance of aquatic invertebrates (e.g., Fig. 1), the diversity in their nutrient-content (Fig. 2), as well as their social, economic and ecological importance (*16*), it is critical to enhance their effective monitoring, assessment and sustainable management in order to steer aquatic food systems towards meeting global, regional and local policy targets. Aquatic invertebrates not only represent an important and diverse source of nutrients, but they also have diverse environmental, social and economic dimensions that also contribute to nutrition security (Fig. 4). They are a nutrient-rich food that are culturally appropriate and accessible in many locations (*41*), with high production potential at relatively low cost (*13*). Policies that enhance their sustainable production and access can tackle multiple environmental and socio-economic targets simultaneously (e.g., “zero hunger”, “gender equality” and “responsible consumption and production”; https://sdgs.un.org/goals).

**Fig. 4.**
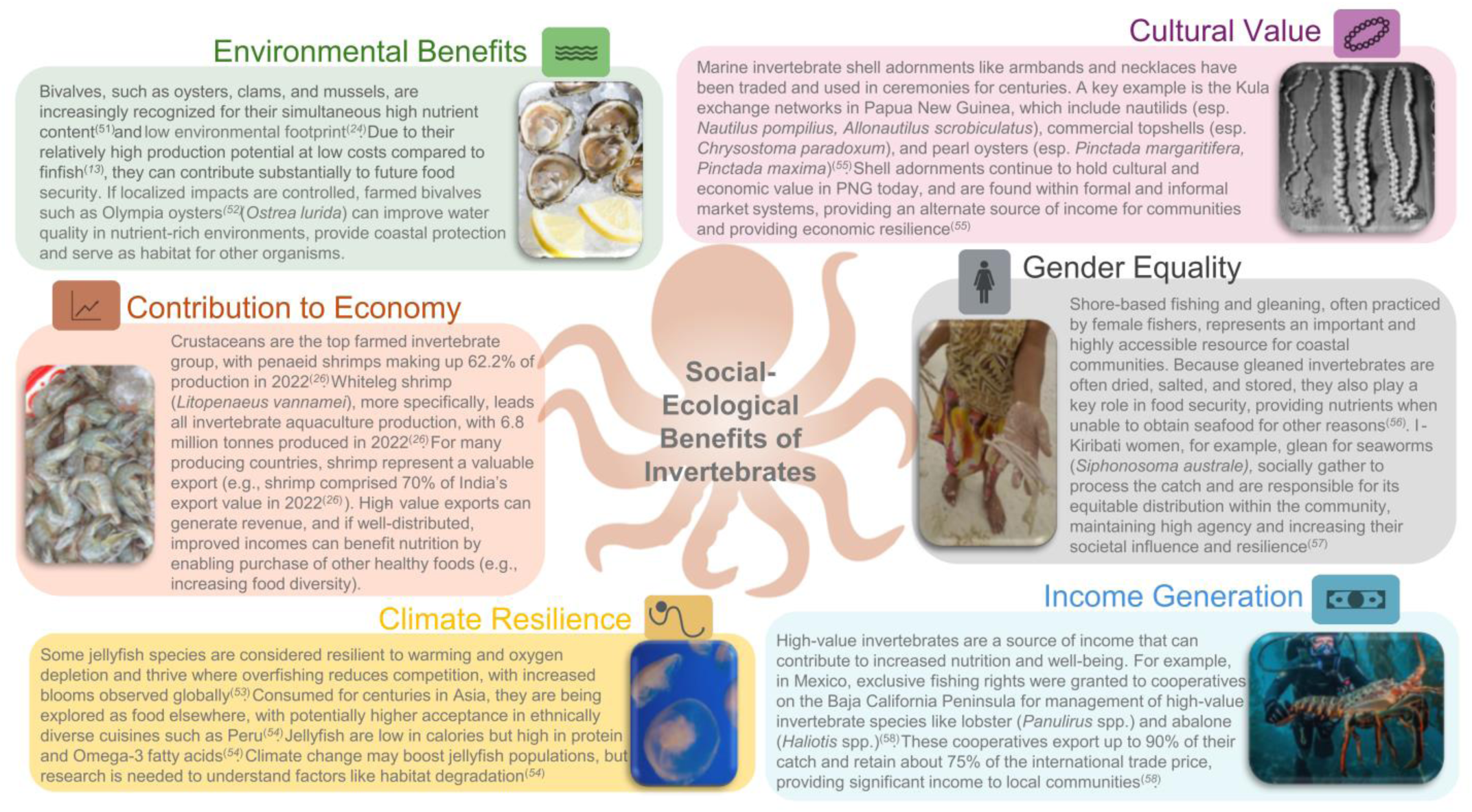
Additional social-ecological benefits of aquatic invertebrates and their indirect contribution to nutrition. Conceptual diagram illustrating the multifaceted roles of aquatic invertebrates, such as bivalves, crustaceans, jellyfish, and sea worms, in providing environmental, economic, cultural and equality-related benefits, in addition to income generation and promoting resilience to climate change. Photo credits: environmental benefit: Taylor Shellfish Farms; contribution to economy: Xufanc; climate resilience: Leslie Wong; cultural value: Adrian Calderon; gender equality: Jacob Eurich; and income generation: Channel Island National Park Authority.

However, overall, their sustainability and climate-resilience is poorly understood (*16*). For example, in terms of catches, we cannot determine if nutrient supplies from invertebrates estimated here (Fig 1) are sustainable because most of the increasingly exploited taxa are unassessed (i.e., we do not know if they are sustainable or not; (*14*)), and those invertebrates that are reported in catches are often grouped into broader taxonomic categories in comparison to vertebrates (e.g., Fig. S12; (*26*)). However, innovative sampling technologies (eDNA (*42*) and automated monitoring (*43*)) in combination with remotely sensed products (e.g., to predict recruitment (*44*)), better accounting of gleaning practices for example in catch monitoring programs (*45*) and traditional field surveys that work under different contexts, such as underwater visual surveys (*46*), mark-recapture (*47*) and community-based mapping methods (*48*), could help increase the resolution and speed of invertebrate data collection, and thus enhance their monitoring, assessment and sustainable management. Invertebrate monitoring is particularly relevant in the context of environmental change (*49*) and increased overfishing (*14*) because it is critical to understand changes that invertebrate aquatic foods are undergoing (*16*) and ensure invertebrate nutrient supplies (e.g., Fig 1) are sustainable and climate-resilient (*50*) as they change.

## Acknowledgments

We thank Angela Zhang, Christina Cornell and Zach Koehn for support in updating and cleaning the Aquatic Food Composition Database, Rashid Sumalia and Daniel Pauly for group discussions and constructive feedback on environmental and life-history traits.

## Funding

Harvard Data Science Initiative (JZM, CDG)

National Science Foundation grant HNDS-I 2121238 (CDG, JAG)

Kenneth K. Chew Endowed Professorship in Aquaculture (JAG)

European Research Council grant 759457 (CCH)

Royal Society University Research Fellowship URF\R1\231087 (JPWR)

## Author contributions

JZM developed and implemented the project with support from CG, CH and JR. JM, KK, DP, and JG were actively involved in project development discussions. NM and SHY cleaned and updated invertebrate specific nutrient composition data with support from JZM and the AFCD development team (see acknowledgements). LA and VP cleaned and updated ecological and environmental trait data for invertebrate species with support from DP. JZM performed the main analyses and wrote the first draft of the manuscript. JGE, LE, JG, WF, and DV wrote individual case studies for Figure 4 with support from JZM. All authors made substantive contributions to the text.

## Data and materials availability

Nutrient composition for aquatic foods (10), marine capture fisheries (Sea Around Us; (25)), invertebrate traits (SeaLifeBase; (27)) and aquaculture production (FishStatJ; (26)) data used for this study are available through online repositories. Updated species-specific invertebrate nutrient composition data used for the predictive models, and all cleaned data and code to replicate the analyses are available for review and will be made publicly available upon publication (i.e., on GitHub and Zenodo). A report of the employed R packages is available in the supplementary information.

## Materials and Methods

### Contribution of aquatic invertebrates to global nutrient supplies

*Global marine capture and aquaculture statistics:* To estimate the contribution of invertebrates to global marine capture fisheries and aquaculture production, we used 2019 reconstructed marine fisheries catches data from the *Sea Around Us* (SAU; (*25*)) and reported aquaculture production from the Food and Agriculture Organization (*26*). The *Sea Around Us* uses officially reported landings from international and national fisheries statistics authorities as a baseline. Reconstructions are then performed by adding estimated unreported catches (e.g., catches discarded at sea or illegal catches) using various literature sources. FishStatJ aquaculture production (*26*) data of territories and land areas were grouped by sovereign nations and reported as alpha-iso 3 country codes (*59*). For instance, all production reported from French Polynesia was included in the production of France (alpha-iso 3 country code ‘FRA’). Aquaculture production species, reported using the Aquatic Sciences and Fisheries Information System (ASFIS) taxonomic reference system in FishStatJ, were converted to scientific species and species groups names (e.g., rainbow trout to *Oncorhynchus mykis*s).

*Assigning nutrient composition data to aquatic foods:* Nutrient concentration estimates per 100g and edible proportions for each individual entry in global data (e.g., fish and invertebrates) were assigned using raw muscle tissue samples within the Aquatic Food Composition Database (AFCD; (*10*)). We selected 29 nutrients important for public health (*11*): 12 minerals (calcium, chromium, copper, iodine, iron, magnesium, manganese, phosphorous, potassium, selenium, sodium and zinc), 11 vitamins (A, C, D, E, B1 (thiamin), B2 (riboflavin), B3 (niacin), B5 (pantothenic acid), B6, B9 (folate) and B12 (cobalamin)), 5 versions of essential fatty acids (total monounsaturated fatty acids (MUFA), total omega 3 fatty acids, total omega 6 fatty acids, eicosapentaenoic acid and docosahexaenoic acid (DHA & EPA) and alpha-linolenic acid (ALA)), and protein. Nutrient composition estimates and edible proportions were assigned hierarchically based on the closest taxonomic resolution (Fig. S9). In other words, for any given nutrient, if a species had species-specific nutrient composition observations in AFCD, we assigned that value. However, if no observed nutrient concentrations were available at the species level, we assigned the mean of the next taxonomic level (e.g., genus). ∼2% and 0.5 % of marine capture yields and aquaculture production weights, respectively, were excluded from the analyses due to a lack of nutrient composition estimates. To obtain nutrient supplies or yields, nutrient concentrations per 100g were multiplied by equivalent units in live weight. Note that we used live weights for our main analyses (i.e., extrapolating the nutrient content from muscle tissue to live weight volumes), but also performed a sensitivity analysis using edible weight (e.g., multiplying live weights by edible proportions; Fig. S1). We grouped nutrient supplies and total catch or production volumes separately for invertebrates and fish, and calculated the percentage of nutrient supplies coming specifically from invertebrates. We separately did this for different sectors, taxa and countries or exclusive economic zones (EEZ). Note that, similar to finfish (*20*), country or EEZ specific nutrient yields are not strongly correlated with the average nutrient concentration of their catches (Fig. S13-S14), indicating, for example, that a country or EEZ with high nutrient yields does not necessarily target the species with most nutrient concentrations.

*Estimating public health relevance of aquatic foods:* Nutrient supplies were converted to the number of yearly requirements met by dividing nutrient-specific nutrient supplies from aquatic invertebrates by their estimated recommended nutrient intakes (RNI), averaged across available demographic groups (e.g., gender and age; Table S3). When available, we used the recommended dietary allowance (RDA; (*60*)). However, when such data was not available we used adequate intake (AI; (*60*)) estimates or values reported in the literature: 433 mg/d for DHA & EPA (i.e., average between different studies reviewed; (*61*)) and 48.8 g/d for monounsaturated fatty acids (i.e., 20% of total energy recommended intake assuming that a gram of MUFA represents 9 kcal; (*62,63*)). Note that, for total omega 6 fatty acids, we used recommended intakes reported for linolenic acids; and for total omega 3 fatty acids, we used the sum of ALA and DHA & EPA recommended intakes (i.e., 1633 mg/d). Yearly nutrient supplies were divided by yearly requirements, ensuring units were the same and assuming 365 days (e.g., yearly requirements in tonnes=daily requirements in tonnes *365). Our approach estimates the number of nutrient requirements met exclusively from the nutrients available from aquatic invertebrates. While we acknowledge that people eat other foods and do not meet their nutritional adequacy exclusively from aquatic invertebrates, this allowed us to compare the importance of aquatic invertebrates across nutrients with a standardized unit relevant for public health.

### Predictive model of invertebrate nutrient concentrations

To estimate the variability and potential drivers of nutrient content in aquatic invertebrates we first updated and validated species-specific invertebrate nutrient concentration samples within AFCD. A total of 16481 samples from 471 invertebrate species were compiled, updated and validated (e.g., reviewing source studies or food composition tables for accuracy, and adding extra information (e.g., sample preparation, relative weight)). Specifically, *food part* (i.e., the body part that was sampled for nutrient concentration) and *food processing* (i.e.,mechanical or chemical processes that transform the animal to the form before consumption) were validated (e.g., examining categorizations for accuracy and disaggregating processing from *sample preparation*; Table S1). Sample sizes and species varied by nutrient (Table S2).

Next, we merged species-specific nutrient data with ecological and environmental trait information available from SeaLifeBase (*27*). Traits were assigned hierarchically, using species-specific data when available or the mean or most common category of the closest taxonomic group (e.g., genus). We included available traits related to energetic demand, thermal regime, habitat and environment that are likely to influence the nutrient composition of aquatic invertebrates (Table S1).

Similar to work done on ray-finned fishes (*20*), we developed a series of Bayesian hierarchical models to predict the nutrient concentration of aquatic invertebrates species based on their taxonomy and ecological and environmental traits, while controlling for other factors thought to impact the nutrient concentration of samples (e.g., food part or processing; Table S1). Note that due to limited species-specific invertebrate sample sizes (Table S2), from the nutrients highlighted, vitamin C and D were not included in this modelling section. Given the structure of our data and the importance of taxonomic identity in explaining the variability in nutrient content (*64*), taxonomy was included in our model hierarchically (i.e., nested random effects of samples within genus, within families, within orders, within class and within phylums). All other factors (e.g., Table S1) were included in the model as fixed effects, with (i) maximum depth log-transformed to normalize the spread of a highly skewed distribution, (ii) all continuous variables standardized by subtracting the mean and dividing by two times the standard deviation (*65*), and (iii) maximum length and length at maturity standardized within the taxonomic level of class (i.e., because for invertebrates, different length types are recorded depending on the taxonomic group). Nutrients were modelled independently due to having different sample sizes (Table S2). For each nutrient, we tested three alternate model structures through leave-out-one cross validation (*66*). That is, a null model including only the intercept, a hierarchical model including only the intercepts reflecting the taxonomic nested structure of our data, and a full model, including taxonomy hierarchically and ecological and environmental traits as covariates. For a minority of nutrients examined (e.g.,vitamin B12), the inclusion of the taxonomic hierarchy alone predicted the out-of-sample data as well as the full model including trait data (Table S4). However, across nutrients (i.e., sum of leave-out-one cross validation information criteria), the full model was significantly preferred in terms of predictive accuracy (Table S4), supporting the inclusion of both taxonomic hierarchy as well as ecological and environmental traits in predicting nutrient concentration for aquatic invertebrates. Thus, for each nutrient, the best linear model structure was:

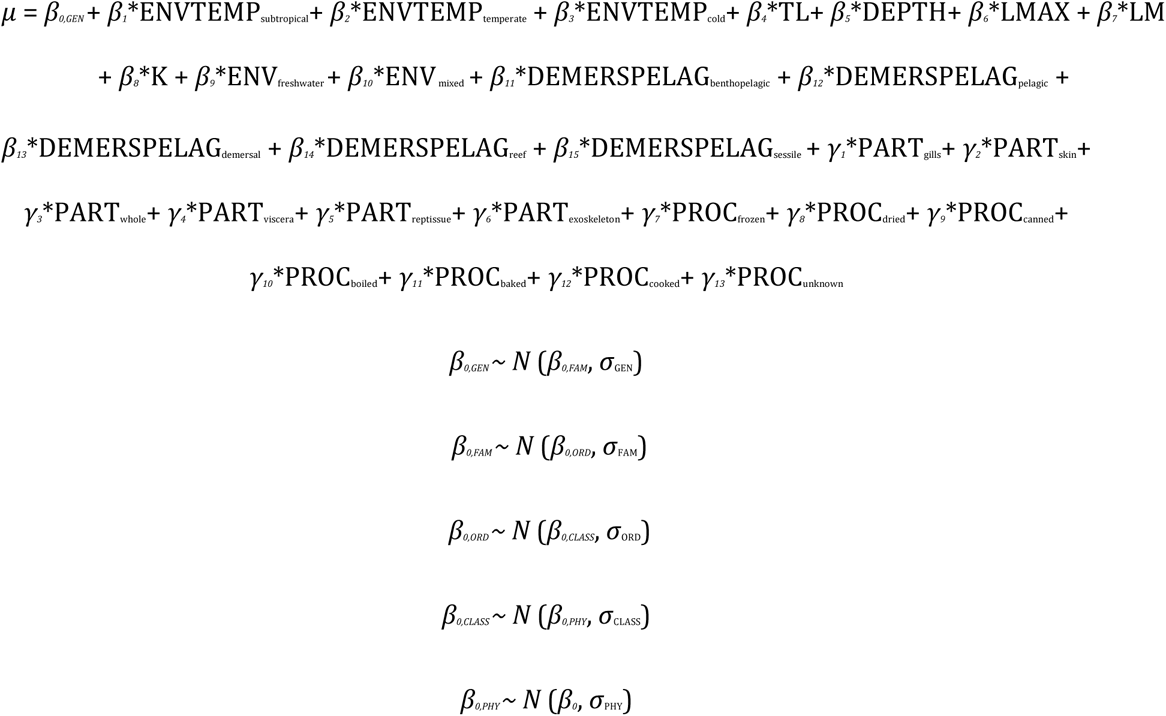

where β0,TAX represents the estimated intercepts at different taxonomic hierarchical levels (GEN: genus, FAM: family; ORD: order, CL: class; and PHY: phylum) for average continuous and most common (i.e., tropical, benthic and marine) covariate categories, β0, is the global estimated intercept, β1-15 represents the estimated parameters for different covariates: thermal regime (ENVTEMP; subtropical, temperate or cold), trophic level (TL), maximum depth (DEPTH), maximum length (LMAX), length at maturity (LM), von Bertalanffy growth parameter (K), environment (ENV; freshwater or mixed) and preferred habitat (DEMERSPELAG; benthopelagic, pelagic, reef-associated, sessile, or demersal), γ1-13 represent the estimated parameters for different nuisance variables: food part (PART; gills, skin, whole or multiple parts, viscera, reproductive tissue, or exoskeleton) and food processing (PROC; frozen, dried, canned, boiled or steamed, baked, grilled or smoked, cooked other, or unknown food prep.); and σTAX represent the estimated standard deviation for the hierarchical taxonomic intercepts.

For each nutrient, we used a hurdle log-normal data likelihood distribution:

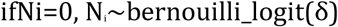

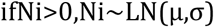

where *Ni* is the nutrient concentration of sample i, δ is the estimated probability of observing a zero for that nutrient, and µ and σ are the mean nutrient concentration informed by the linear model structure above and the standard deviation, both of the variable’s natural logarithm.

Models were run in Rstan (*67*) through the brms package (*68*) and using default priors. Four chains were run for each scenario, leaving 4000 samples in the posterior distribution of each parameter. Convergence was monitored by running four chains from different starting points, examining posterior chains and distribution for stability, checking that the potential scale reduction factor (also termed R_hat) was close to 1 (below 1.01) and examining the effective sample sizes (>400). Identifiability was examined by inspecting posteriors vs. prior distributions and by calculating posterior contraction values (*69*). We examined posterior predictive distributions to check for model fit. Additionally, we checked model residual patterns against variables not included in our model due to high missingness (e.g., Fig. S15-S16). All final models reported converged, fit well the data and had high posterior contraction (e.g., Fig. S17-S18).

### Predicting nutrient composition of invertebrates in SeaLifeBase

Finally, we used model parameters estimated in the above section to predict nutrient composition of all macroinvertebrates available in SeaLifeBase (*27*). The initial database contained a total of 71,591 species. From these, a total of 55,864 species were identified as macroinvertebrates based on taxonomic groupings: 281 were confirmed as “edible” based on literature (e.g., (*70–75*)) and 50,807 were identified as potentially edible (excluding 834 non-edible species and 4,223 species from the phyla *Bryozoa, Platyhelminthes, Porifera and Priapulida*, which we assumed were not edible). Based on available literature (*25*, *27, 70, 71*) some potentially edible macroinvertebrate species (1,083) were also categorized based on their “fisheries importance” into the following groups: industrial, highly commercial, commercial, minor commercial, subsistence fisheries and bycatch. We provide nutrient composition estimates for all invertebrate species; but, in our analyses, we filtered the database to include only species classified as potentially edible macroinvertebrates (e.g., Fig.3). Note the initial database also included ascidians (phylum Chordata) which are not invertebrates. However, they were included in model predictions given (i) similarities in some ecological and environmental traits with invertebrates, and (ii) that some are edible (*76*).

Some species had missing information. However, to perform predictions for all available species we took some assumptions. For example, if a specific-group (e.g., Cnidaria) did not have a group-specific estimated intercept (i.e., we did not have that group for that nutrient when building the Bayesian hierarchical models), we used the upper level intercept (e.g., global intercept) in combination with ecological and trait information. Similarly, if a species was missing some ecological or environmental trait information, we assumed the baseline category (e.g., mean for continuous variables and most common categories: benthic, marine, and tropical). To represent the data visually, we divided predicted nutrient concentrations by per capita daily recommended nutrient intakes (see above), restricting values to one when they were above such value (i.e., if 100g of muscle tissue provided >100% of recommended nutrient intakes).

Ultimately, we used matched predictive nutrient concentrations in terms of recommended nutrient intakes of all potentially-edible macroinvertebrate species to species-specific occurrence data (*34*) to show the geographical public health potential of macroinvertebrates (i.e., what nutrient concentrations from invertebrates may be available in different geographical contexts).

### Case studies

The author group developed six case studies representing a diverse array of geographies, target species, spatial scales and management contexts to highlight the additional socio-ecological benefits of aquatic invertebrates and their indirect contribution to nutrition (Fig. 4).

**Fig. S1.**
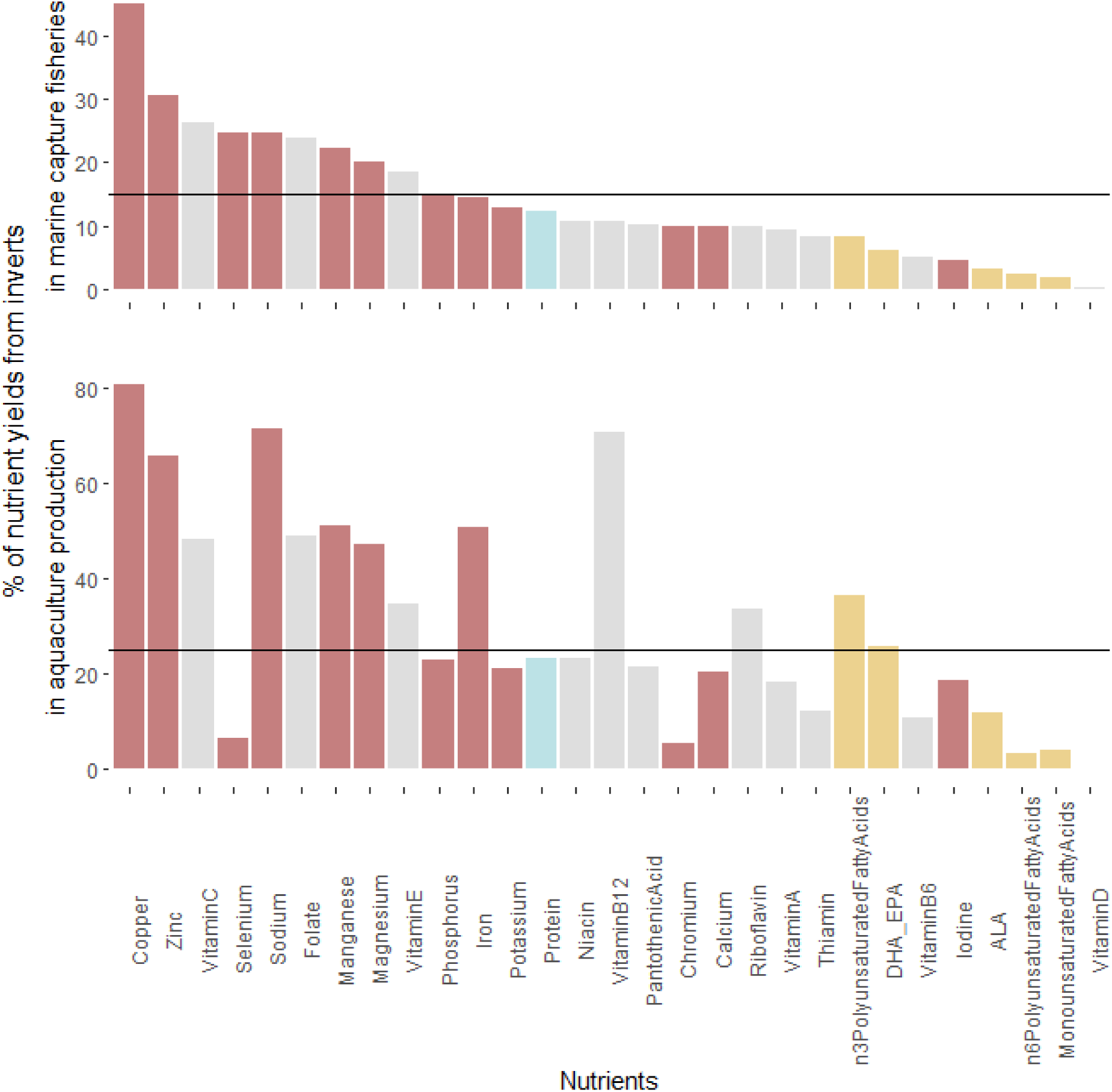
Nutrient contribution of aquatic invertebrates to marine capture fisheries and aquaculture production using edible weights instead of live weights (i.e., Fig. 1). Bars are color-coded by nutrient type (i.e., minerals, vitamins, protein or fatty acids). Horizontal line indicates the contribution of invertebrates to total catch and aquaculture production volumes, respectively.

**Fig. S2.**
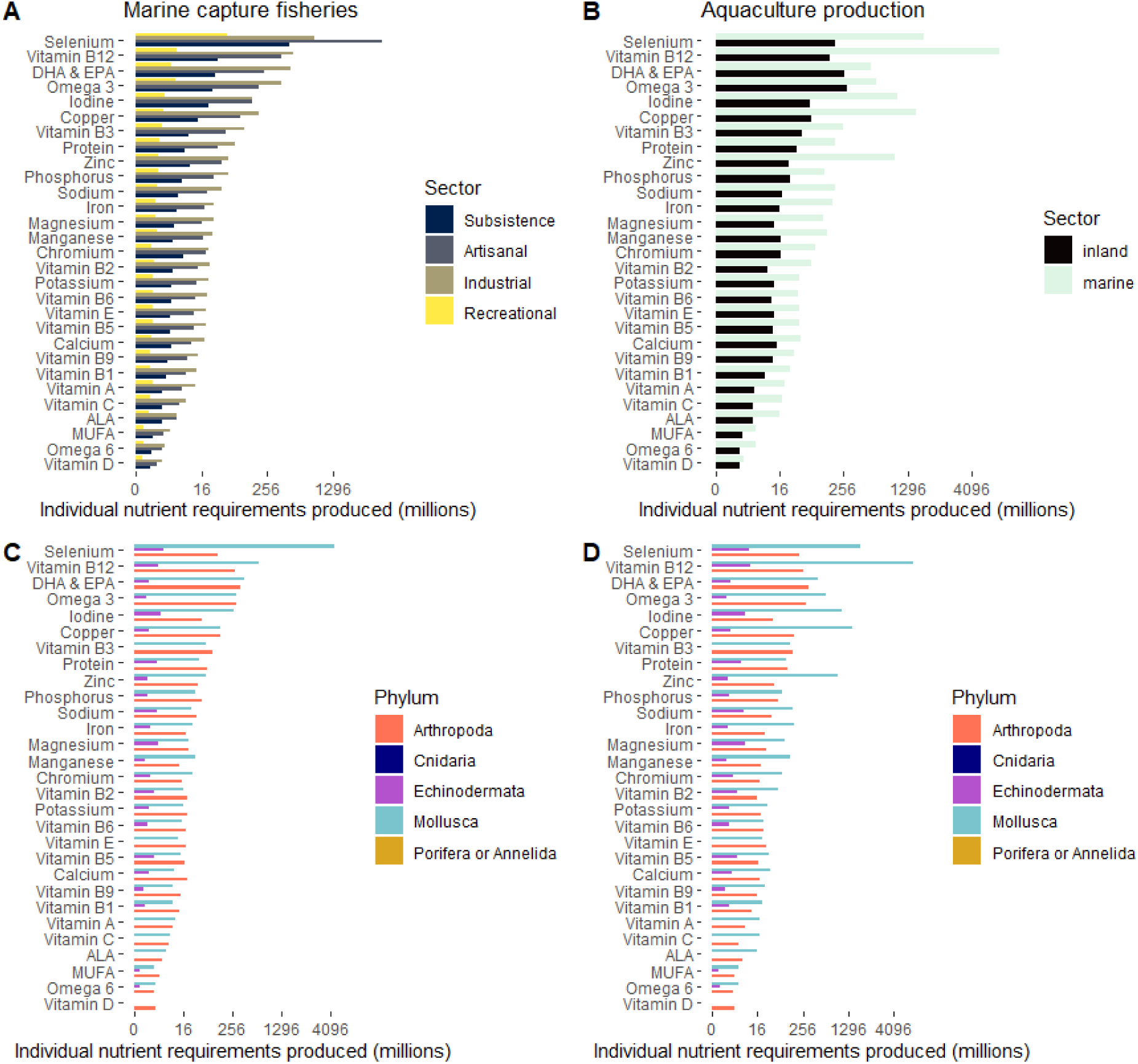
Nutrient contribution of aquatic invertebrates separated by sector and phyla. Number (in millions) of yearly nutritional requirements produced from aquatic invertebrates in marine capture fisheries (**A,C**) or aquaculture production (**B,D**) for 2019. For each nutrient, we use the average requirements across demographic groups (Table S3). To aid clarity the x axes were fourth-root transformed and reported values are on an arithmetic scale.

**Fig. S3.**
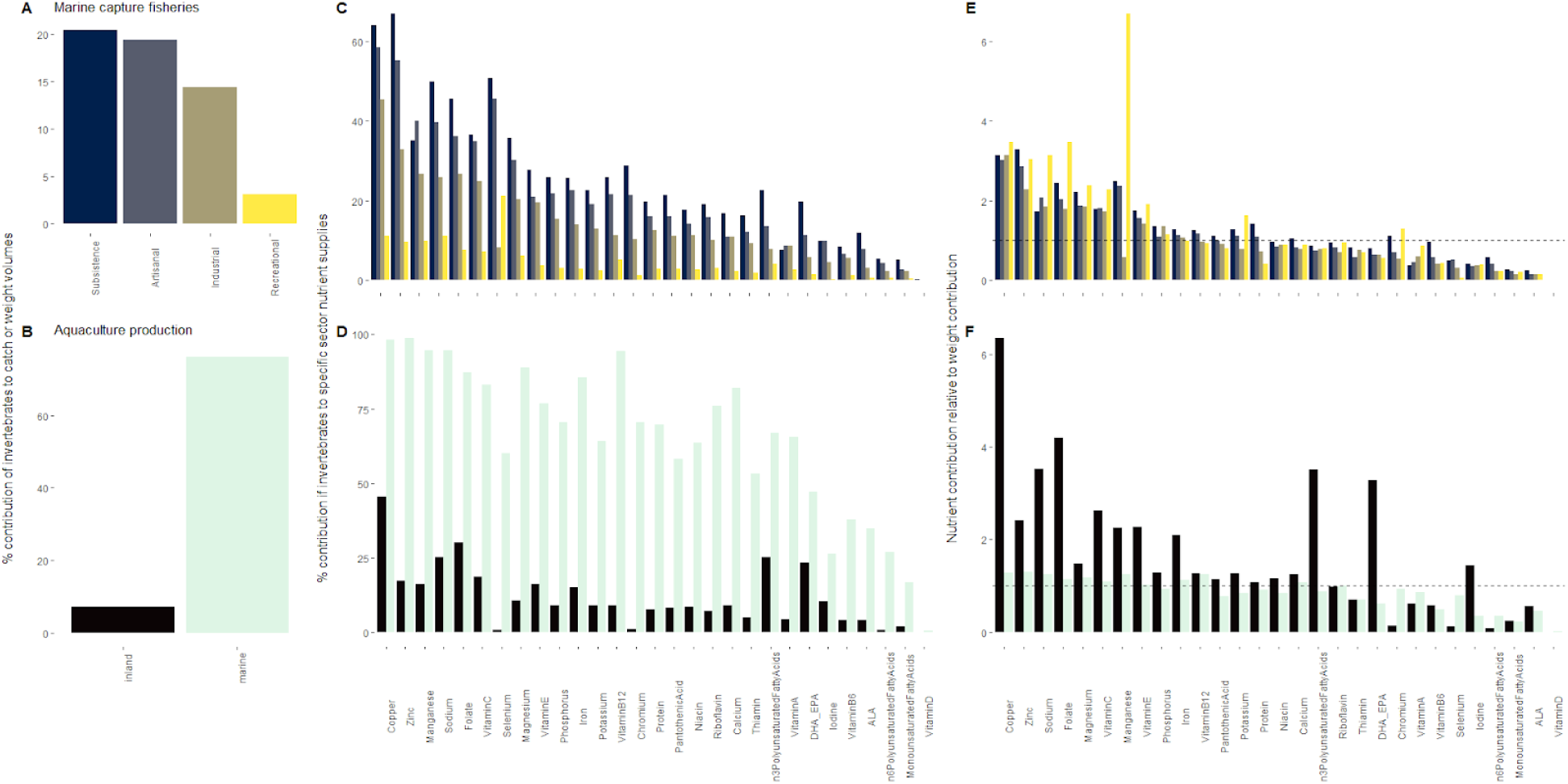
Nutrient contribution of aquatic invertebrates to marine capture fisheries and aquaculture production separated by sectors. Bars are color-coded by sector. **(A-B)** % contribution of invertebrates to marine catch **(A)** or aquaculture production **(B)** volumes (i.e., weight). **(C-D)** % contribution of invertebrates to nutrient supplies in marine capture fisheries **(C)** and aquaculture production **(D)**. **(E-F)** Nutrient contribution relative to catch or weight contribution for marine capture fisheries **(E)** or aquaculture production **(F)**. Horizontal lines in **E-F** indicate when weight contributions are equal to nutrient contributions (i.e., bars above the line indicate invertebrates contribute disproportionately to that nutrient for that sector).

**Fig. S4.**
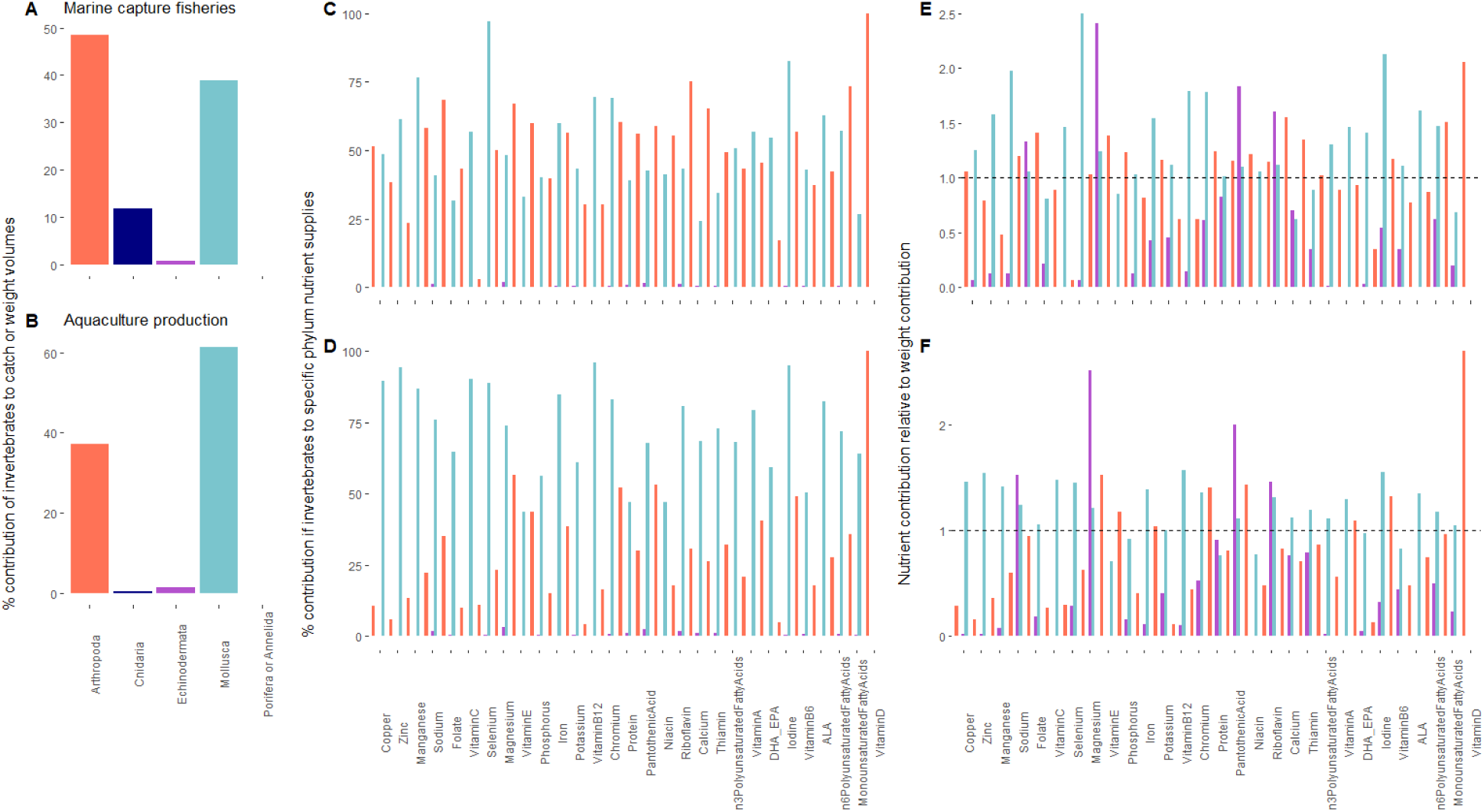
Nutrient contribution of aquatic invertebrates to marine capture fisheries and aquaculture production separated by phylum. Bars are color-coded by phylum. **(A-B)** % contribution of invertebrates to marine catch **(A)** or aquaculture production **(B)** volumes (i.e., weight). **(C-D)** % contribution of invertebrates to nutrient supplies in marine capture fisheries **(C)** and aquaculture production **(D)**. **(E-F)** Nutrient contribution relative to catch or weight contribution for marine capture fisheries **(E)** or aquaculture production **(F)**. Horizontal lines in **E-F** indicate when weight contributions are equal to nutrient contributions (i.e., bars above the line indicate invertebrates contribute disproportionately to that nutrient for that phylum).

**Fig. S5.**
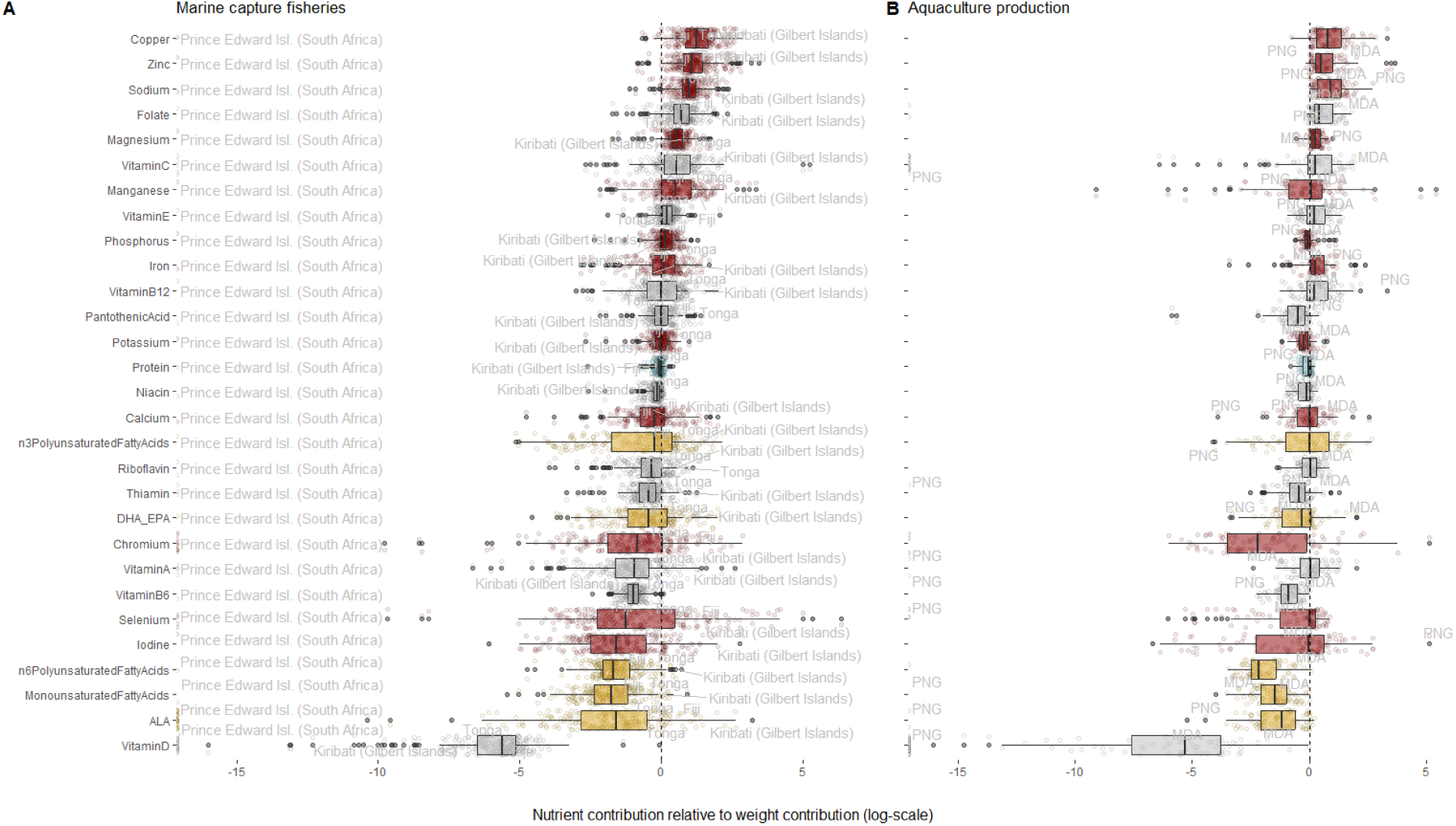
Country specific variability in nutrient contribution of aquatic invertebrates relative to country-specific volume contributions for (A) marine capture fisheries and (B) aquaculture production (e.g., log(% contribution of invertebrates to copper supplies/ % contribution of invertebrates to catch volumes)). Each point is an EEZ **(A)** or country (ISO3c; **B**) and grey labels are the EEZ or ISO3c that ranked first or last (i.e., highest or lowest) in terms of overall nutrient contributions from invertebrates relative to weight volumes. The vertical line indicates when nutrient contribution and catch contribution of invertebrates is the same (i.e., points to the right of the line indicate invertebrates contribute disproportionately to that nutrient for those countries). Boxplots and points are coloured-coded by nutrient type (i.e., minerals, vitamins, protein or fatty acids). In **(B)** MDA and PNG refer to Moldova and Papua New Guinea, respectively.

**Fig. S6.**
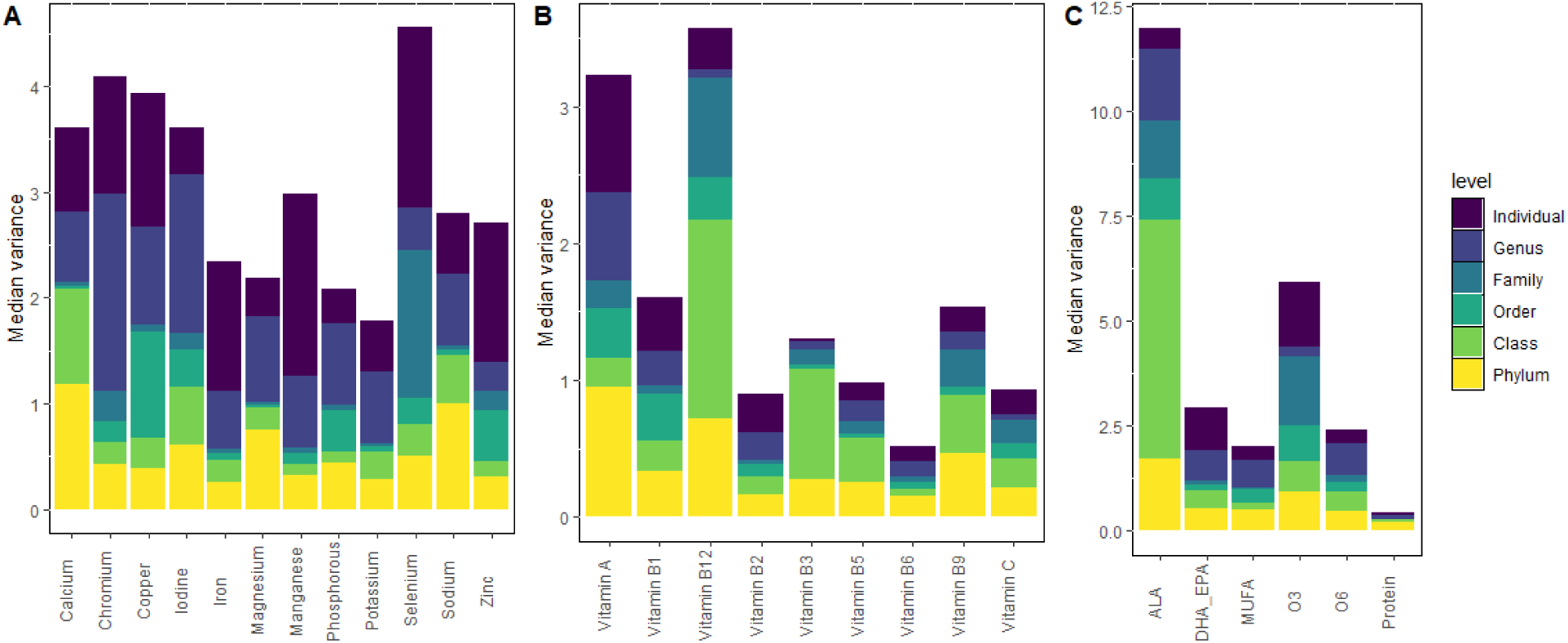
Estimated variance decomposition of our Bayesian Hierarchical models predicting nutrient content in aquatic invertebrates. Each column is a nutrient for which models were performed. Estimated variance for each group (nested) is stacked and color-coded by the hierarchical level.

**Fig. S7.**
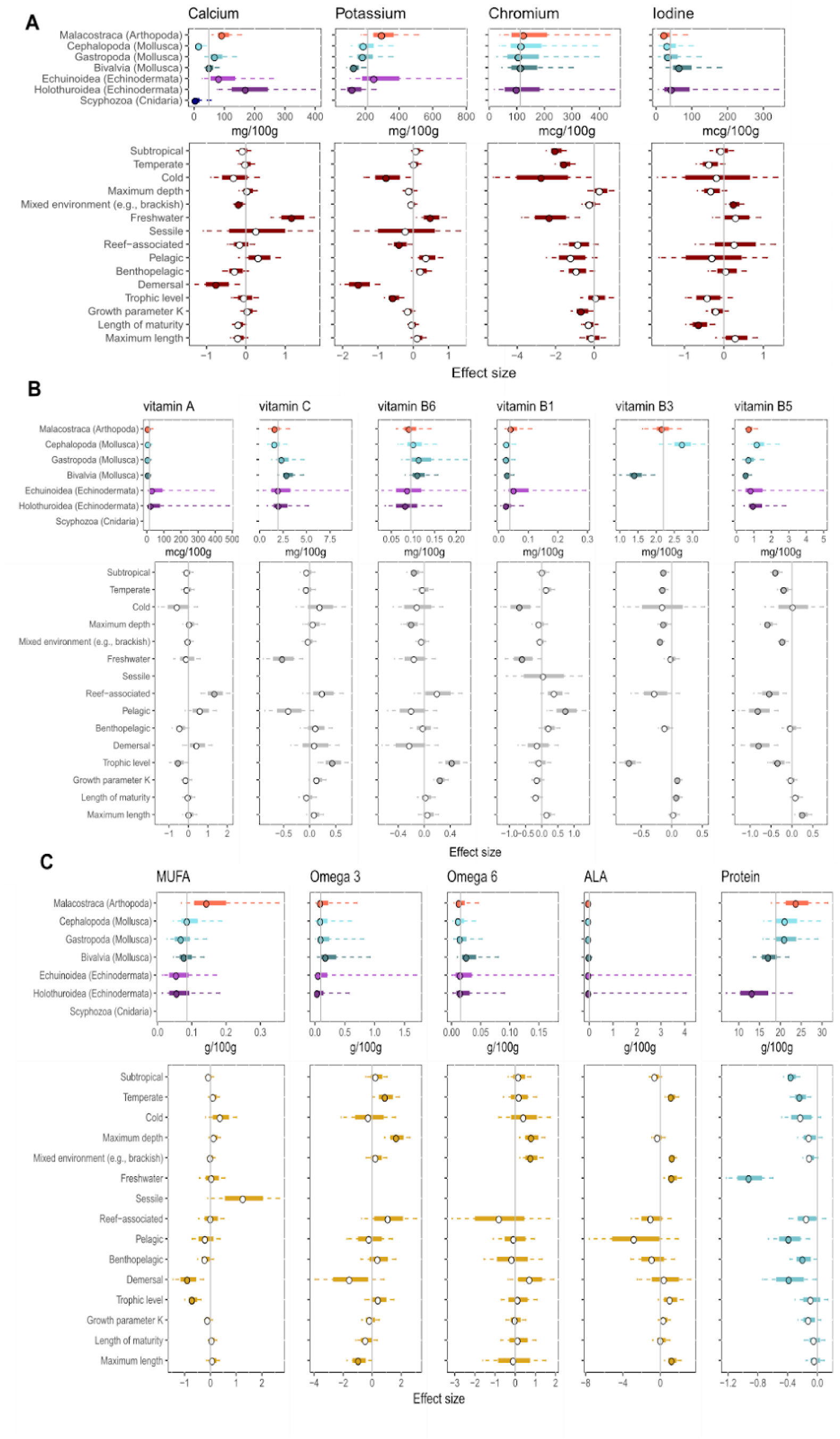
Variability and potential drivers of nutrient content in aquatic invertebrates for nutrients not highlighted in Fig. 2. **(A)** minerals, **(B)** vitamins and **(C)** macronutrients: abbreviations represent MUFA (Monounsaturated fatty acids), DHA_EPA (eicosapentaenoic acid and docosahexaenoic acid), O3FA (omega 3 fatty acids), O6FA (omega 6 fatty acids) and ALA (alpha-linolenic acid). As in Fig. 2, top rows are the estimated nutrient concentration per taxonomic class. Points are the median estimated nutrient concentration for that class (i.e., global intercept plus relevant phylum and class intercept offsets), solid and dashed lines represent 50 and 90% uncertainty intervals, respectively. Vertical line represents the median estimated global intercept. Results are colour-coded by class. Bottom rows are the effect sizes of different ecological and environmental factors on average nutrient concentrations (in log scale). Points are medians, solid and dashed lines represent 50 and 90 % uncertainty intervals, respectively. Vertical lines are set at 0 (i.e., the baseline category). Nutrients are colour-coded by nutrient type. Estimates are baselined for raw muscle tissue at average environmental and ecological continuous conditions (e.g., depth, maximum length, length of maturity, growth parameter K and trophic level) and most common categories (e.g., tropical, marine, and benthic).

**Fig. S8.**
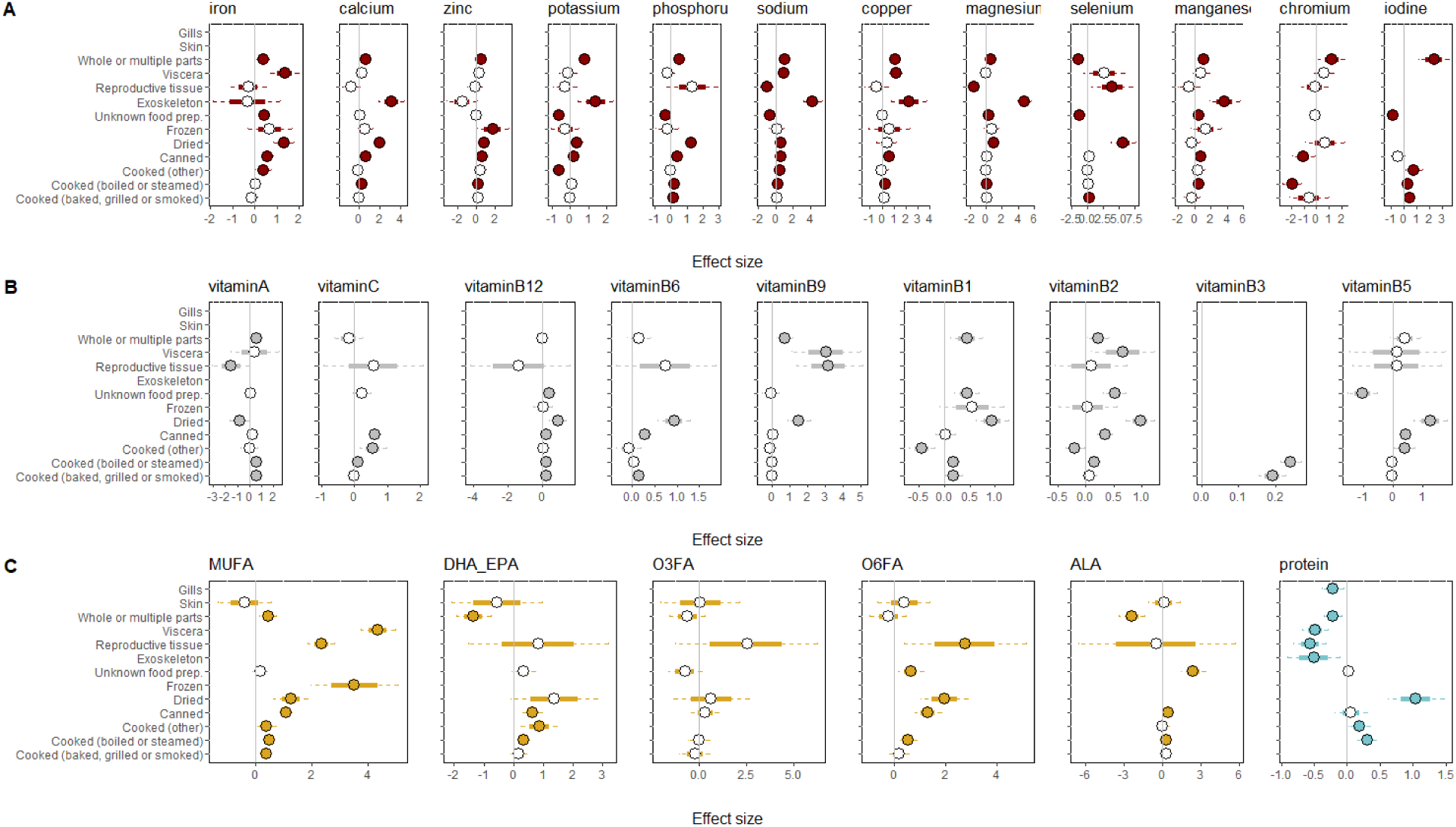
Effect size of nuisance covariates accounted for in our model. Each plot is a nutrient color-coded by nutrient type, separated by **(A)** minerals, **(B)** vitamins and **(C)** macronutrients: abbreviations represent MUFA (Monounsaturated fatty acids), DHA_EPA (eicosapentaenoic acid and docosahexaenoic acid), O3FA (omega 3 fatty acids), O6FA (omega 6 fatty acids) and ALA (alpha-linolenic acid). Points are medians, thick lines represent 50% uncertainty intervals and dashed lines represent 90% uncertainty intervals. If the parameter 90% uncertainty intervals overlapped zero, the point is not filled. Estimates are baselined for raw muscle tissue at average environmental and ecological continuous conditions (e.g., depth, Lmax, Lm, K, TL) and most common categories (e.g., tropical, marine, and benthic).

**Fig. S9.**
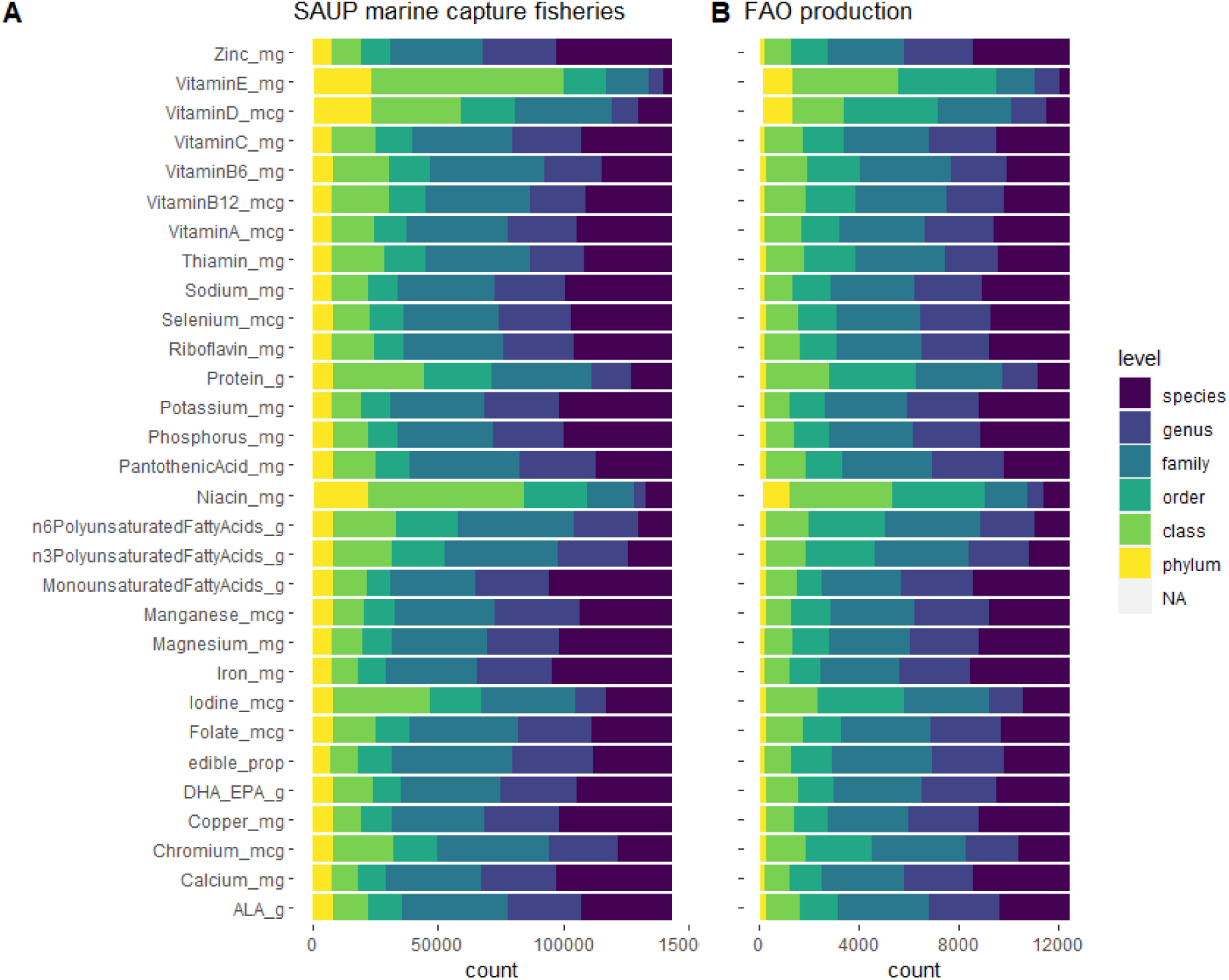
Taxonomic resolution of observed nutrient concentration and edible proportion data. Taxonomic level of assigned observed nutrient or edible proportion data from the Aquatic Food Composition Database (AFCD) for marine capture fisheries **(A)** and aquaculture production **(B)** data used to calculate nutrient contributions of aquatic invertebrates.

**Fig. S10.**
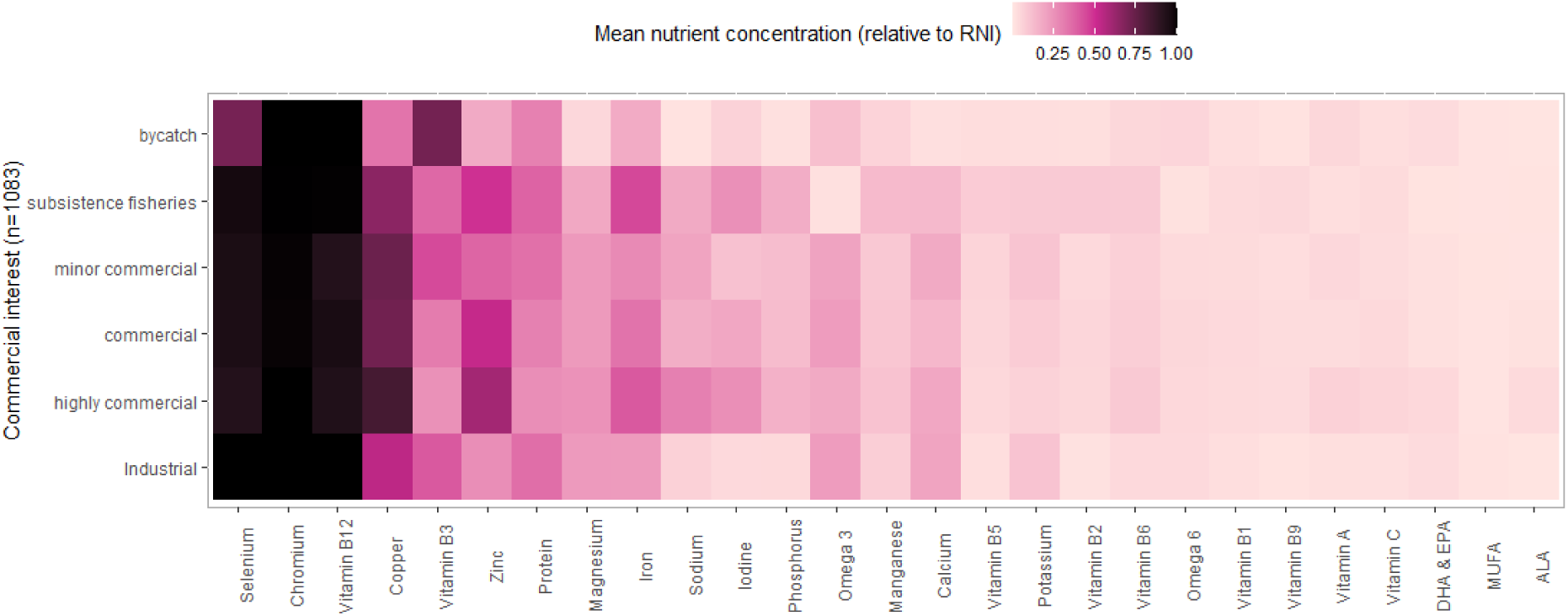
Nutrient predictions relative to recommended nutritional intakes (RNI) for 1,083 potentially edible macroinvertebrates available in SeaLifeBase (*27*) categorized as of fisheries importance. Tiles are color-coded by the mean nutrient concentration of the species in a given group relative to RNI.

**Fig. S11.**
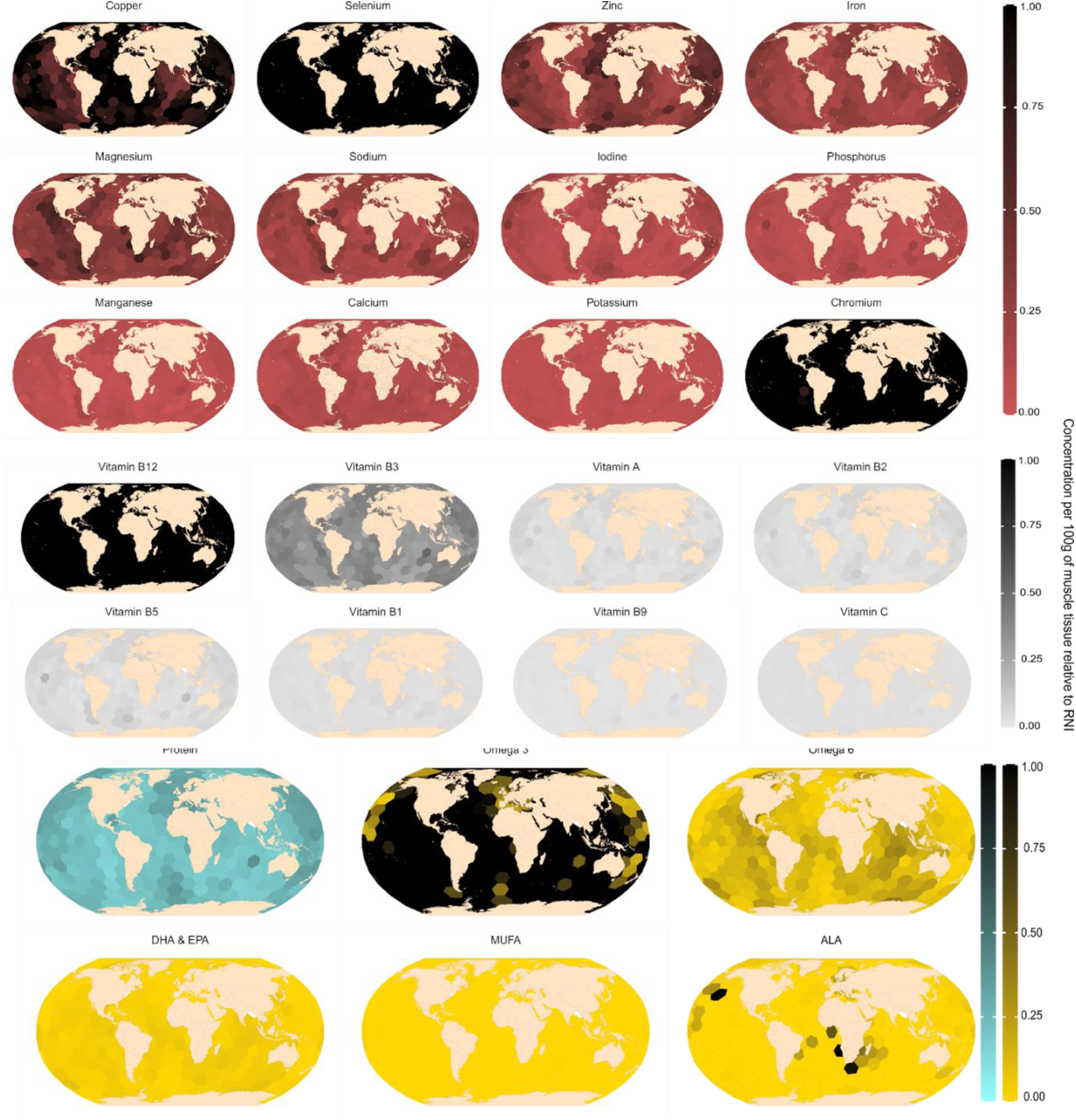
Geographic distribution of potentially edible aquatic macroinvertebrates based on species occurrences and estimated nutrient content. Each plot is an individual nutrient represented as the nutrient concentration per 100g of muscle tissue relative to RNI: minerals **(A)**, vitamins **(B)** and protein and fatty acids **(C)**. Values above one, were restricted to 1.

**Fig. S12.**
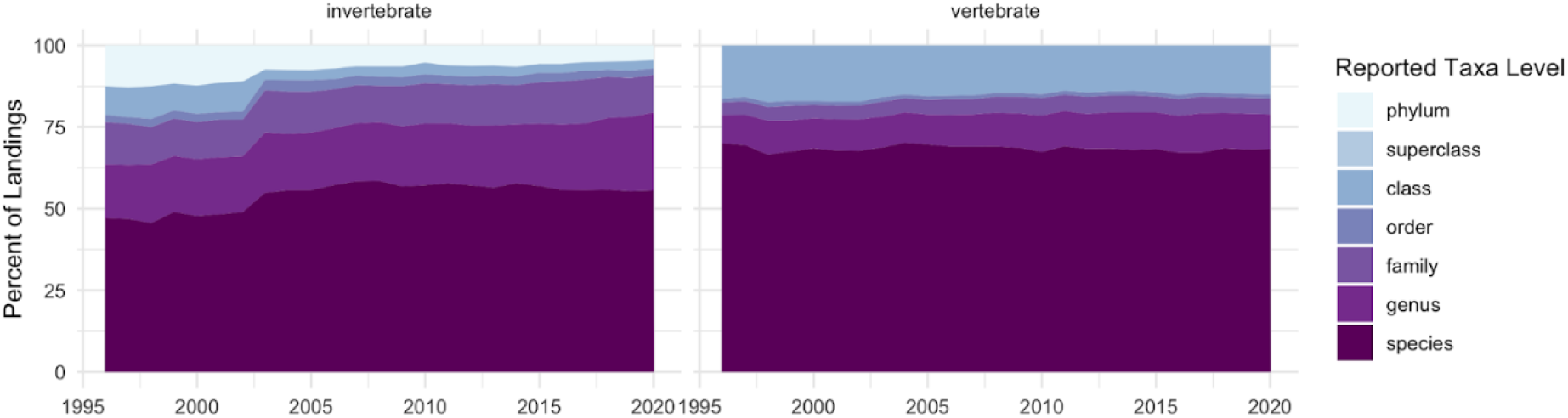
Taxonomic resolution of reported invertebrate catches in comparison to vertebrates, represented as percent of landings through time (*26*,*57*).

**Fig. S13.**
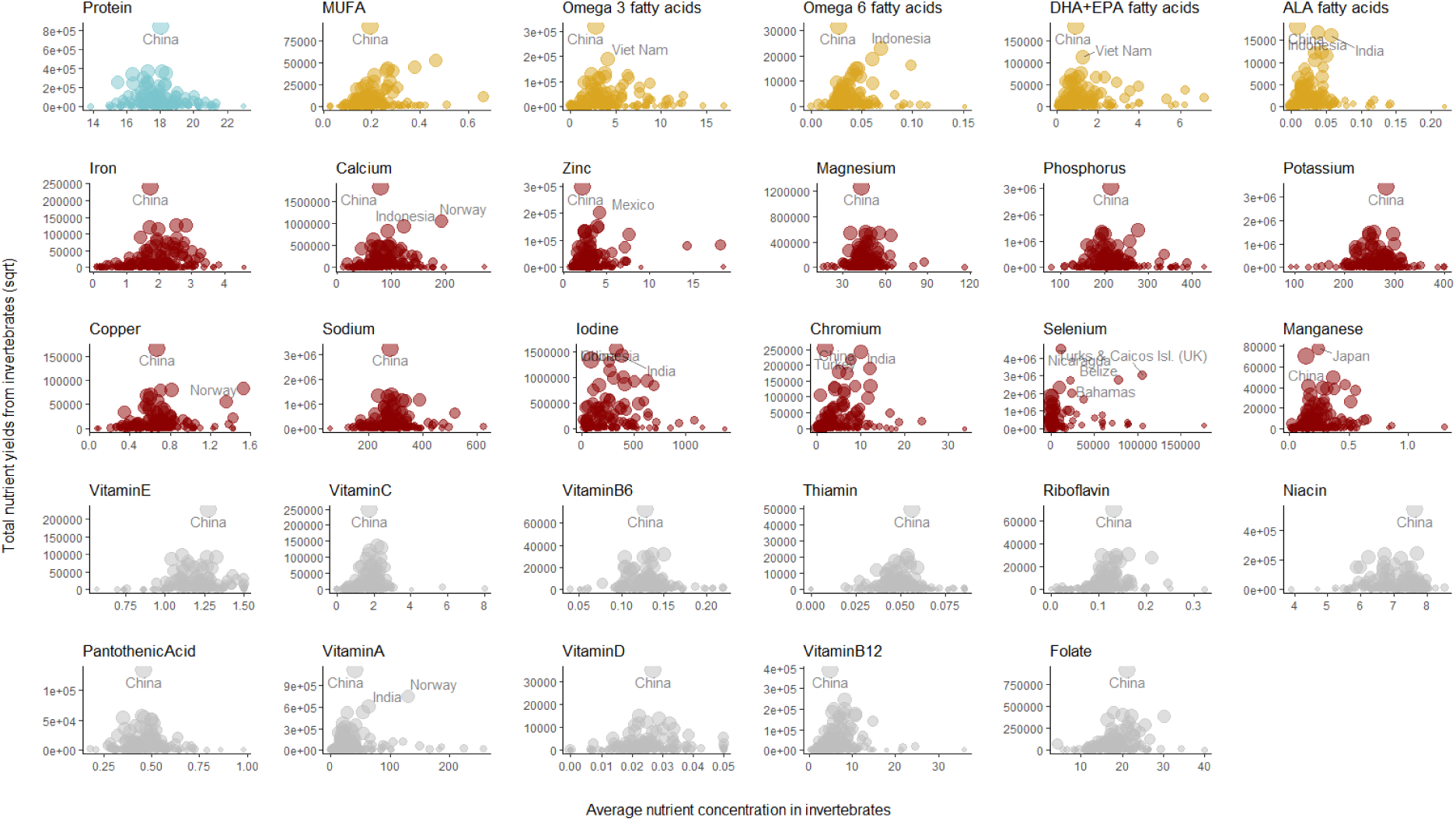
Relationships between invertebrate nutrient yield (sqrt transformed), concentration and total catch for each nutrient examined. Each point is a fishing entity and size is scaled according to total invertebrate catch (also sqrt transformed).

**Fig. S14.**
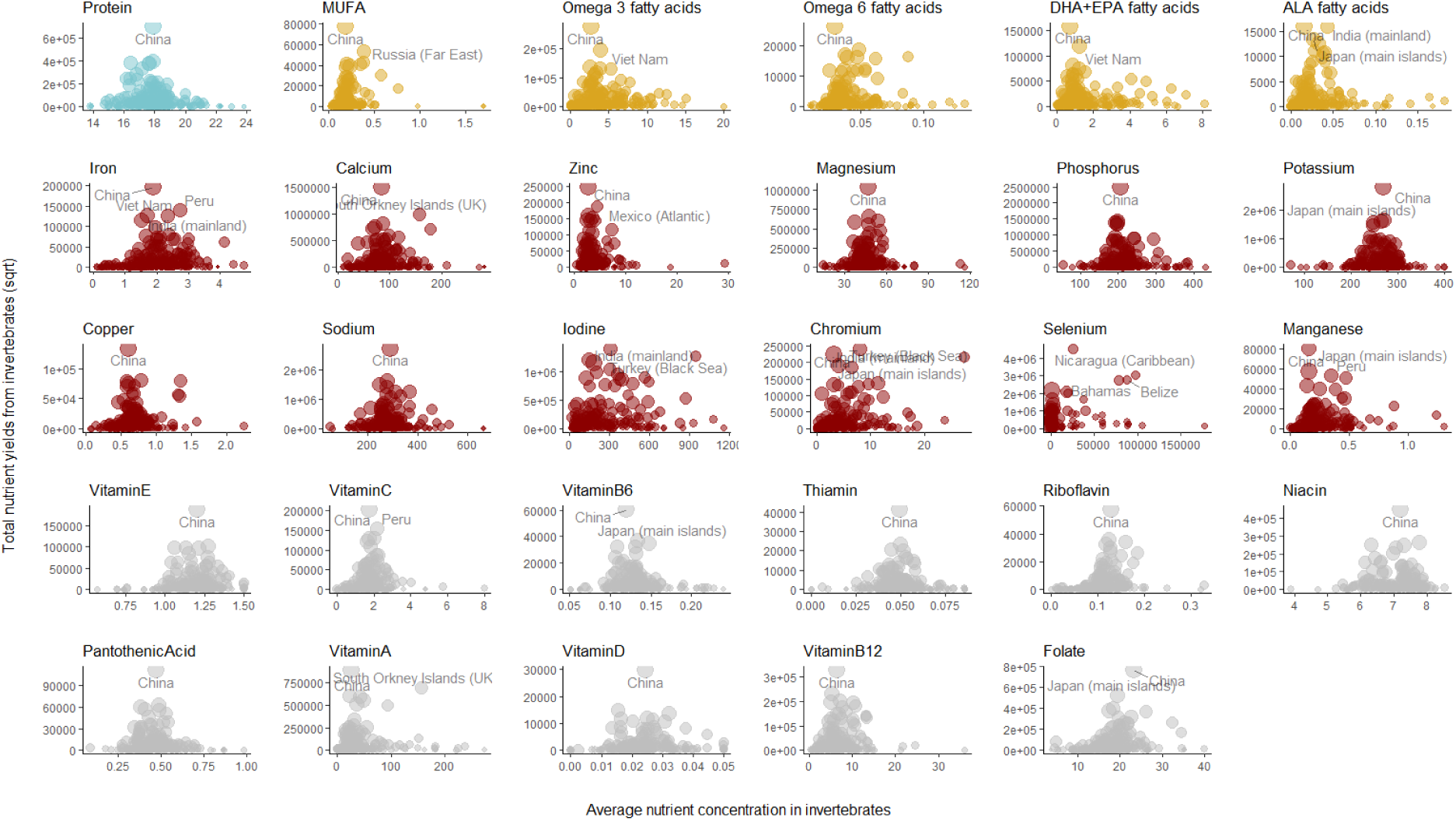
Relationships between invertebrate nutrient yield (sqrt transformed), concentration and total catch for each nutrient examined. Each point is an EEZ and size is scaled according to total invertebrate catch (also sqrt transformed).

**Fig. S15.**
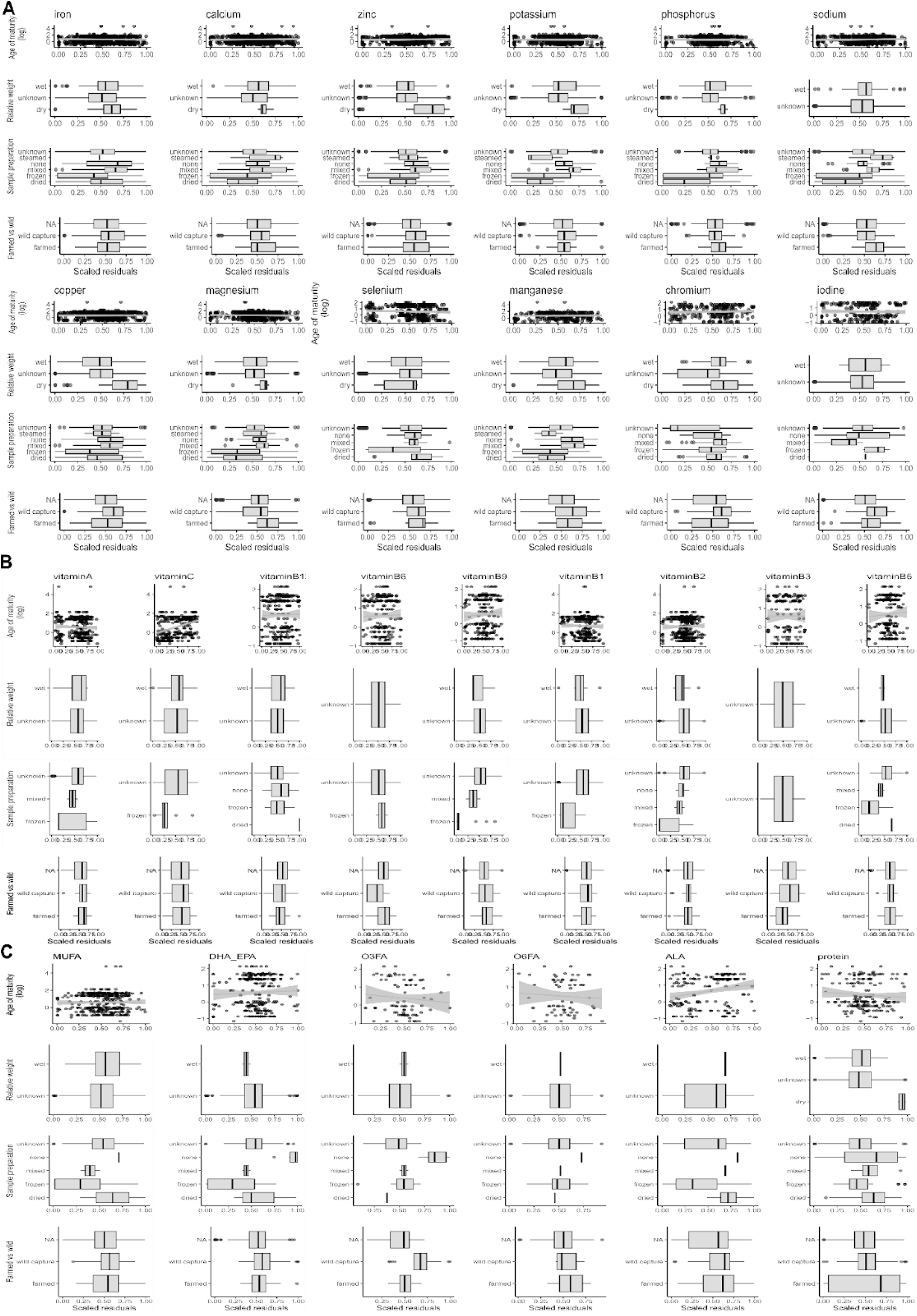
Residual patterns (scaled residuals) with omitted covariates (age of maturity, relative weight, sample preparation and farmed vs. wild) due to data missingness. **(A)** minerals, **(B)** vitamins and **(C)** macronutrients: abbreviations represent MUFA (Monounsaturated fatty acids), DHA_EPA (eicosapentaenoic acid and docosahexaenoic acid), O3FA (omega 3 fatty acids), O6FA (omega 6 fatty acids) and ALA (alpha-linolenic acid).

**Fig. S16.**
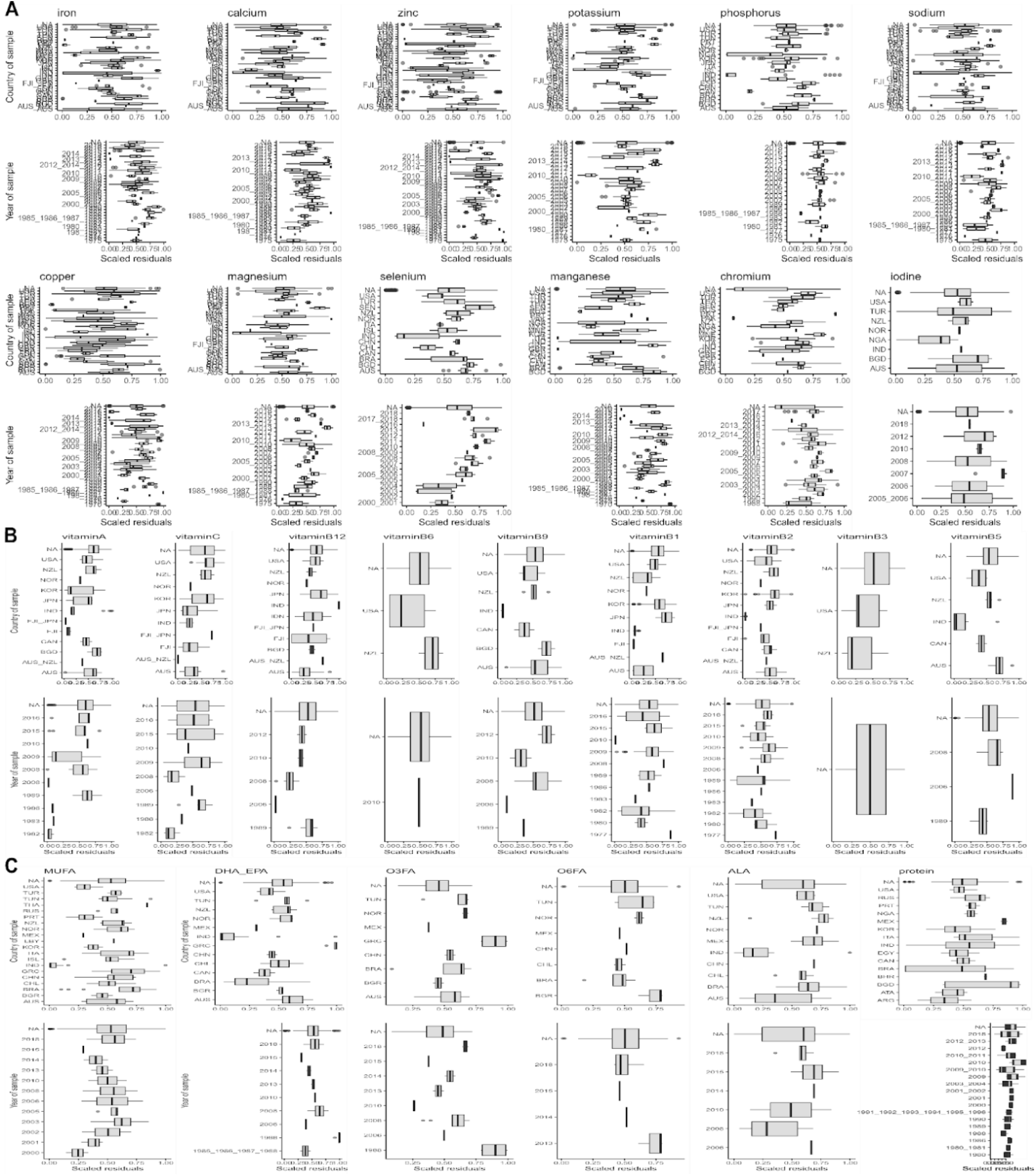
Residual patterns (scaled residuals) with omitted covariates (year of sampling and country of sampling) due to data missingness. **(A)** minerals, **(B)** vitamins and **(C)** macronutrients: abbreviations represent MUFA (Monounsaturated fatty acids), DHA_EPA (eicosapentaenoic acid and docosahexaenoic acid), O3FA (omega 3 fatty acids), O6FA (omega 6 fatty acids) and ALA (alpha-linolenic acid).

**Fig. S17.**
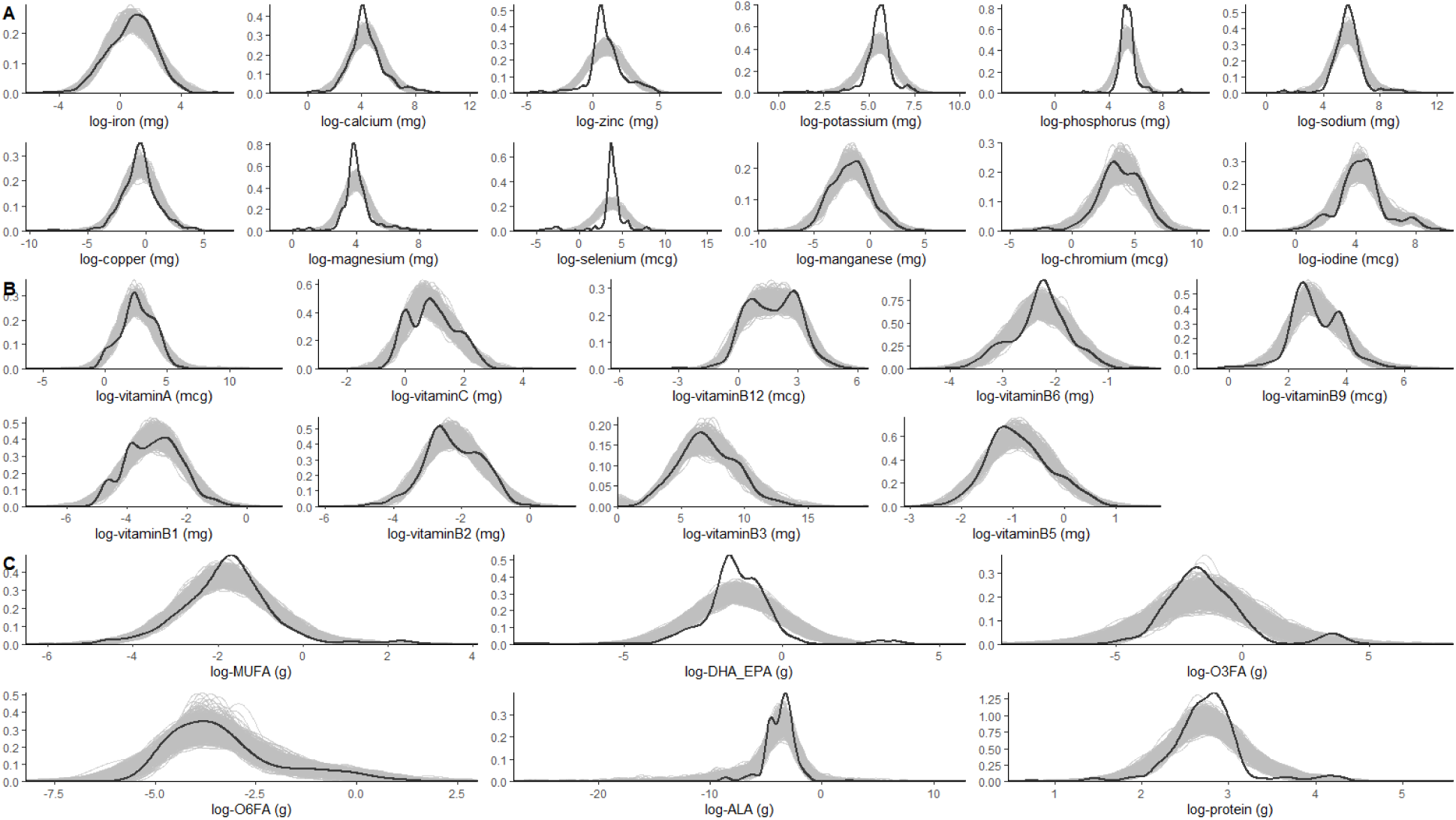
Model fits of our Bayesian Hierarchical Models. Density of posterior draws (grey) vs observed data (black). Each plot is a nutrient (concentration in log-scale), separated by **(A)** minerals, **(B)** vitamins and **(C)** macronutrients: abbreviations represent MUFA (Monounsaturated fatty acids), DHA_EPA (eicosapentaenoic acid and docosahexaenoic acid), O3FA (omega 3 fatty acids), O6FA (omega 6 fatty acids) and ALA (alpha-linolenic acid).

**Fig. S18.**
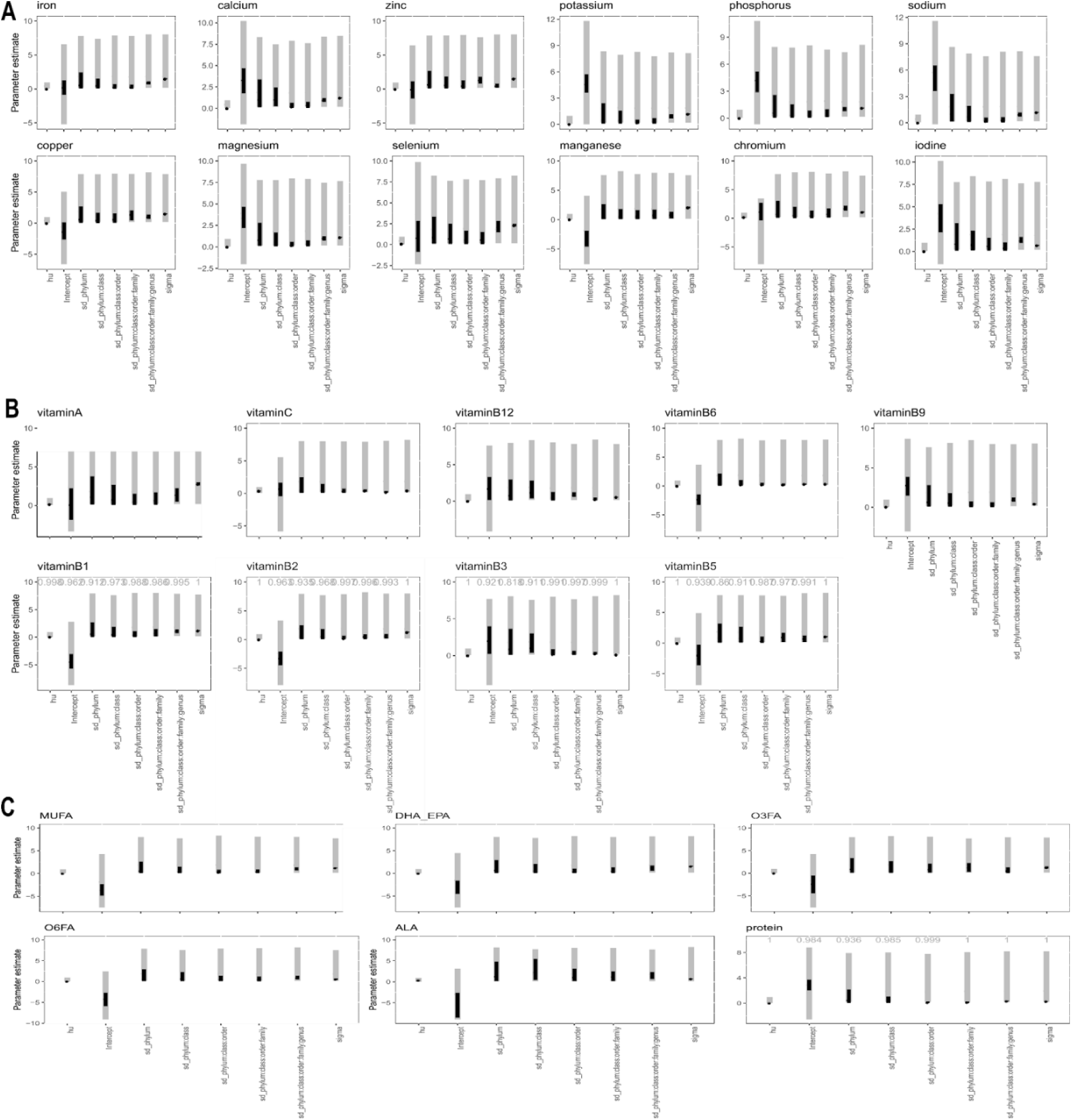
Identifiability checks. Prior (grey) vs posteriors (black). **(A)** minerals, **(B)** vitamins and **(C)** macronutrients: abbreviations represent MUFA (Monounsaturated fatty acids), DHA_EPA (eicosapentaenoic acid and docosahexaenoic acid), O3FA (omega 3 fatty acids), O6FA (omega 6 fatty acids) and ALA (alpha-linolenic acid).

**Table S1.**
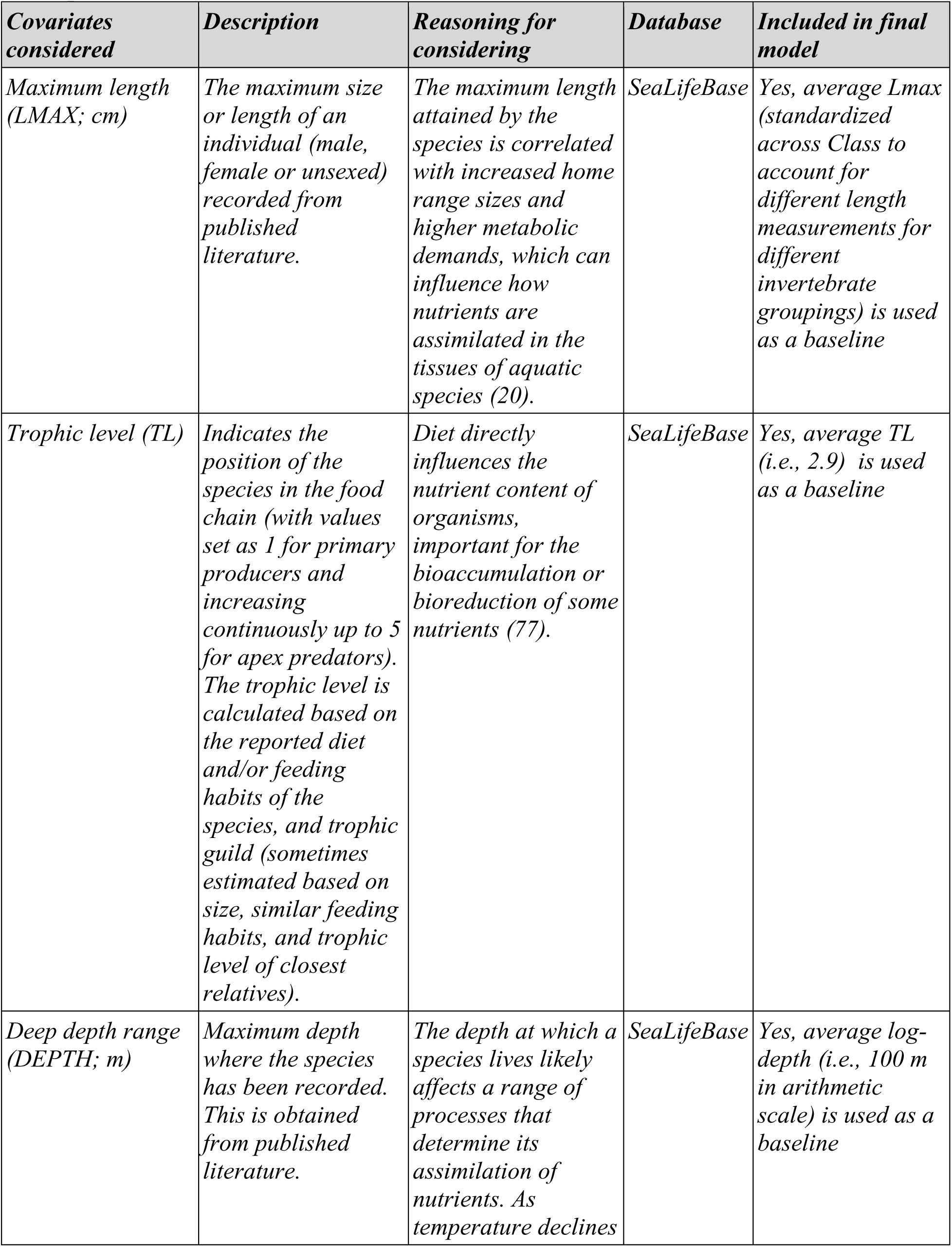
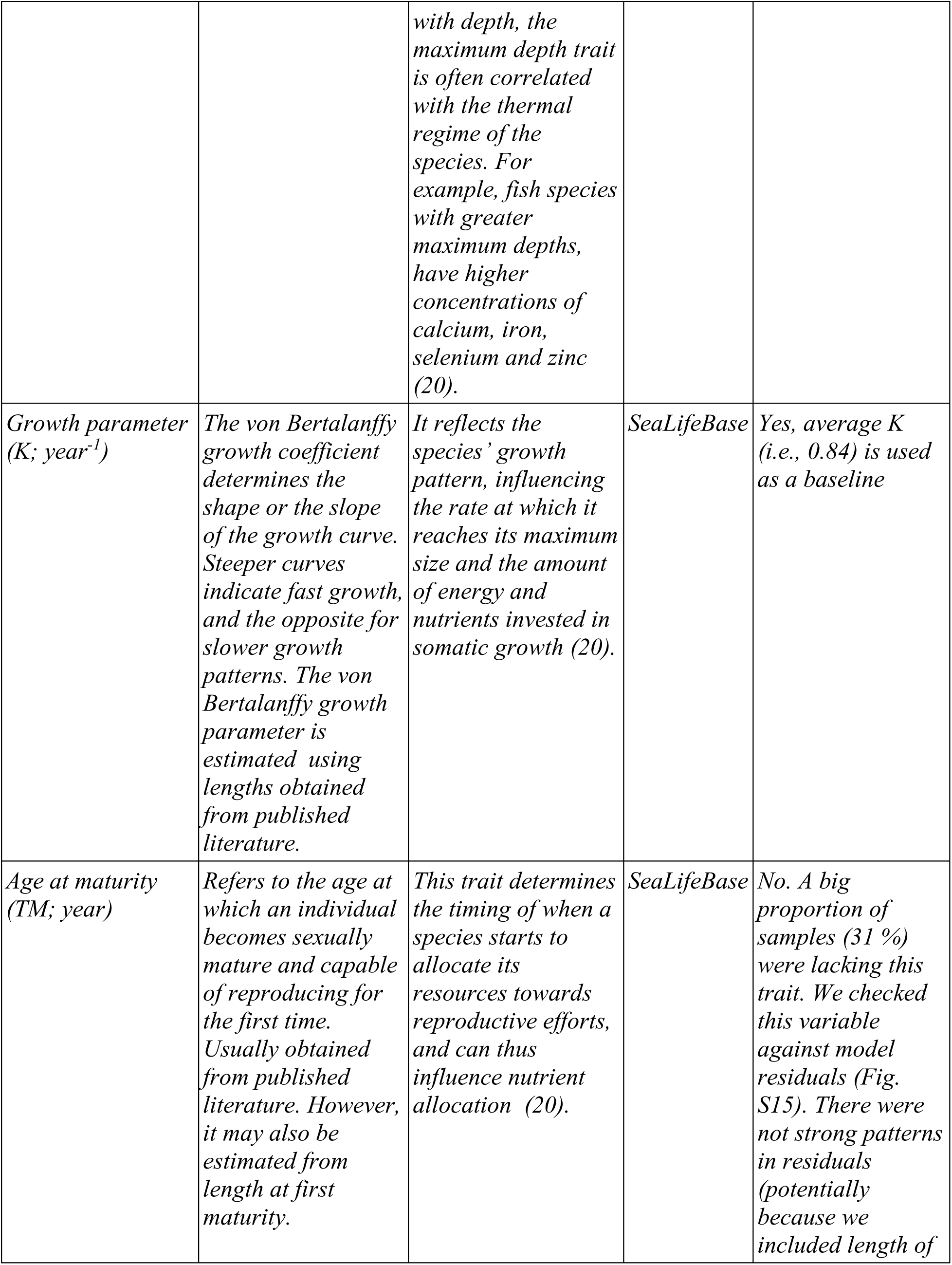
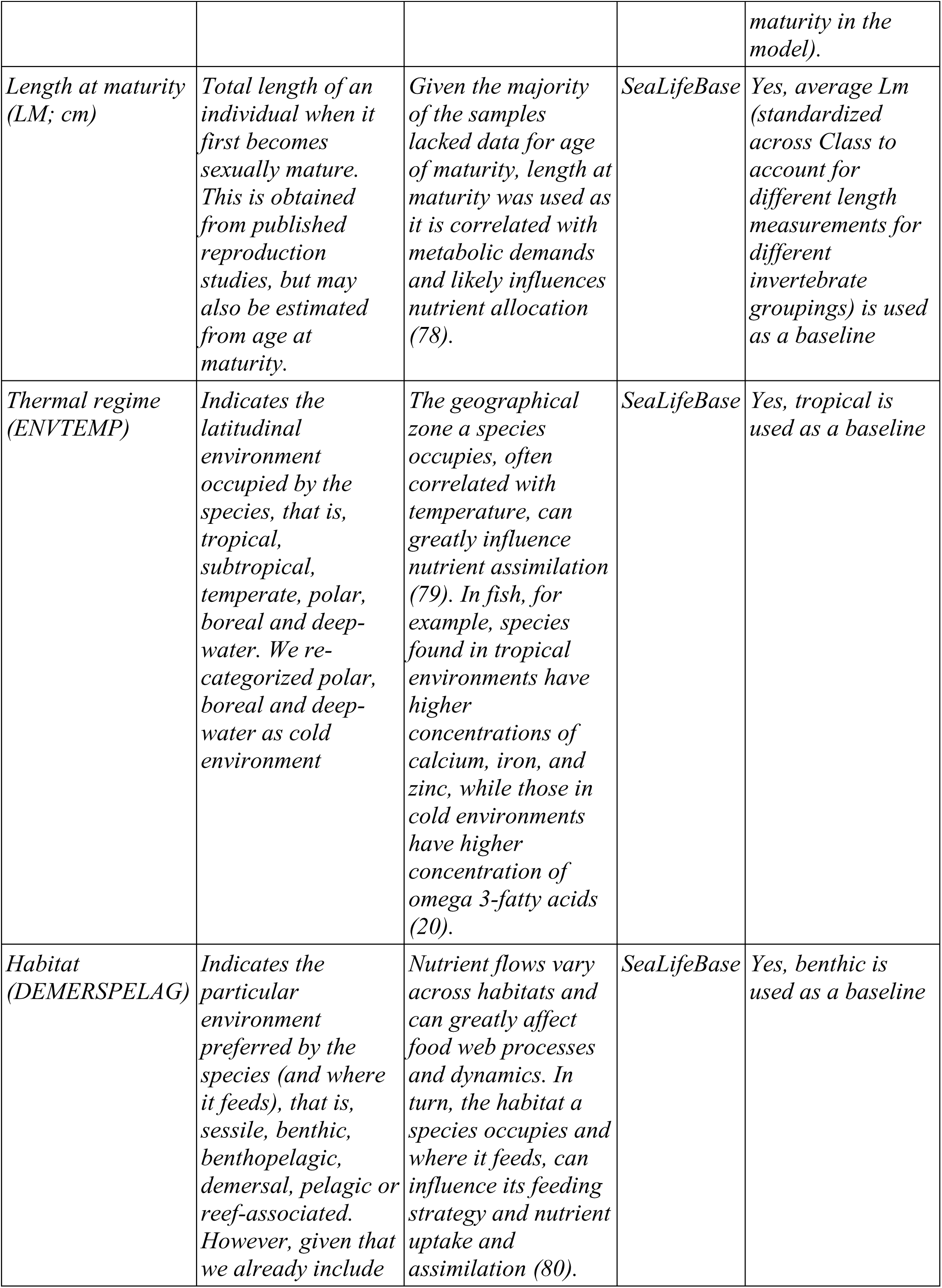
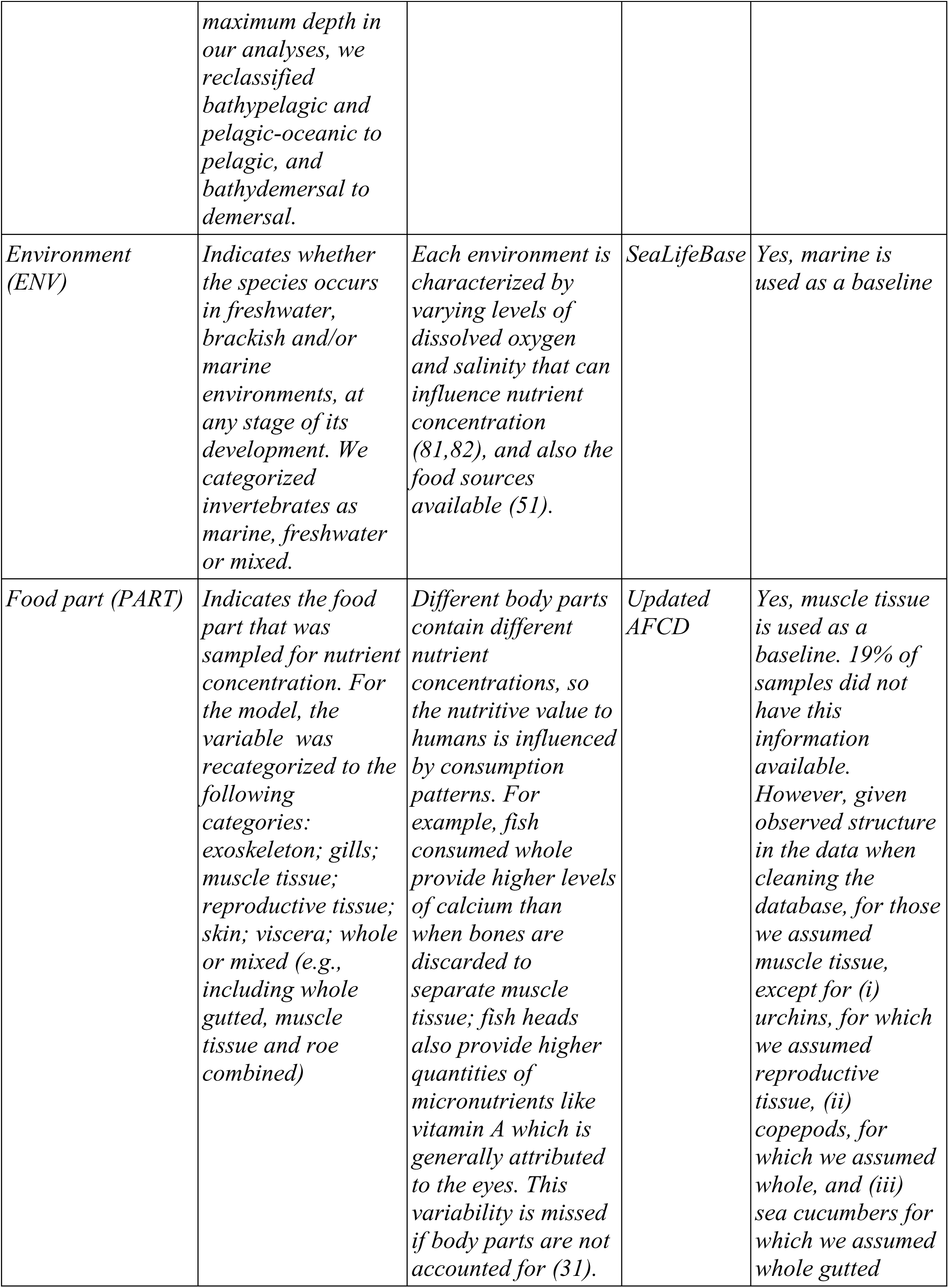
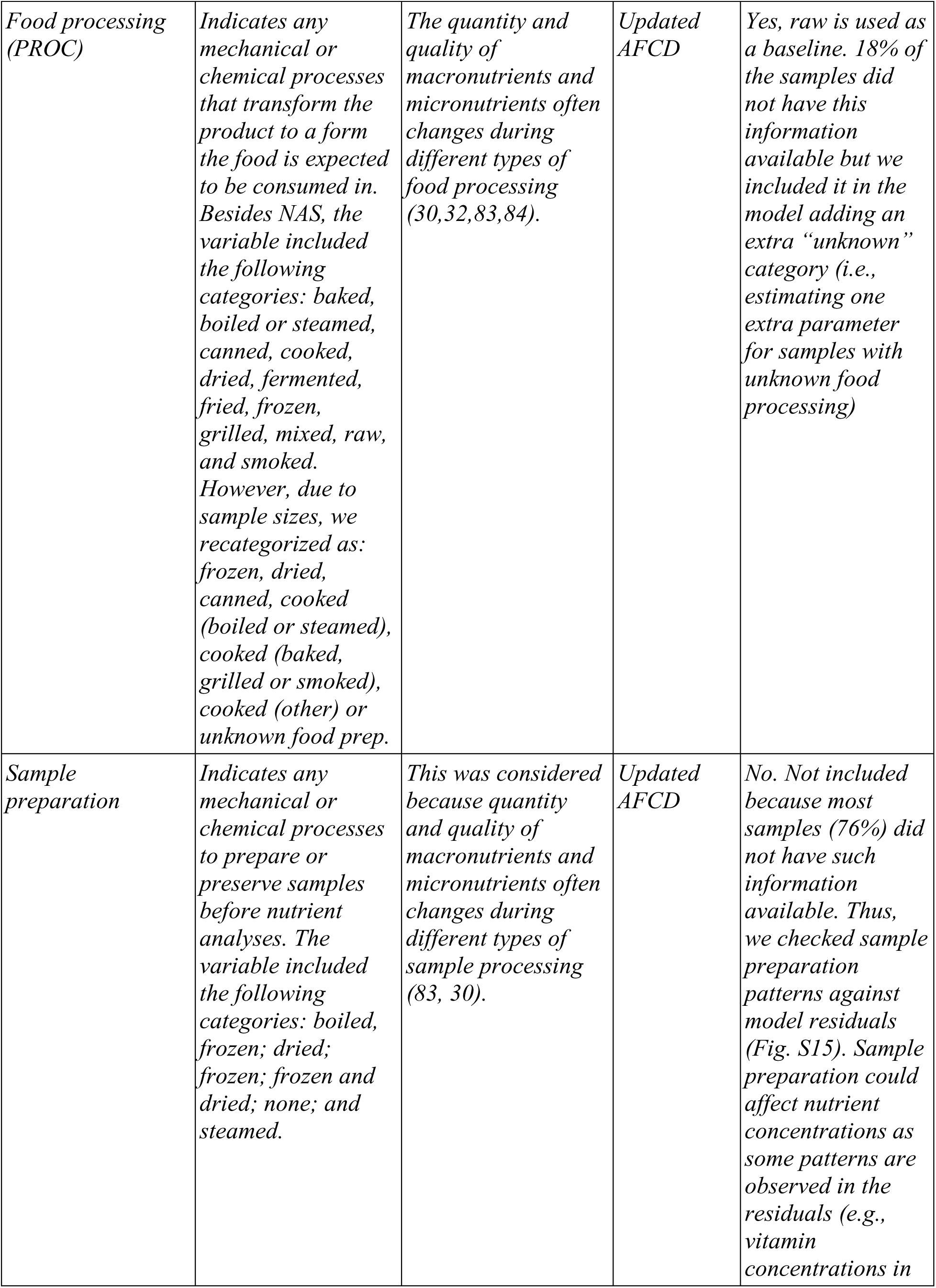
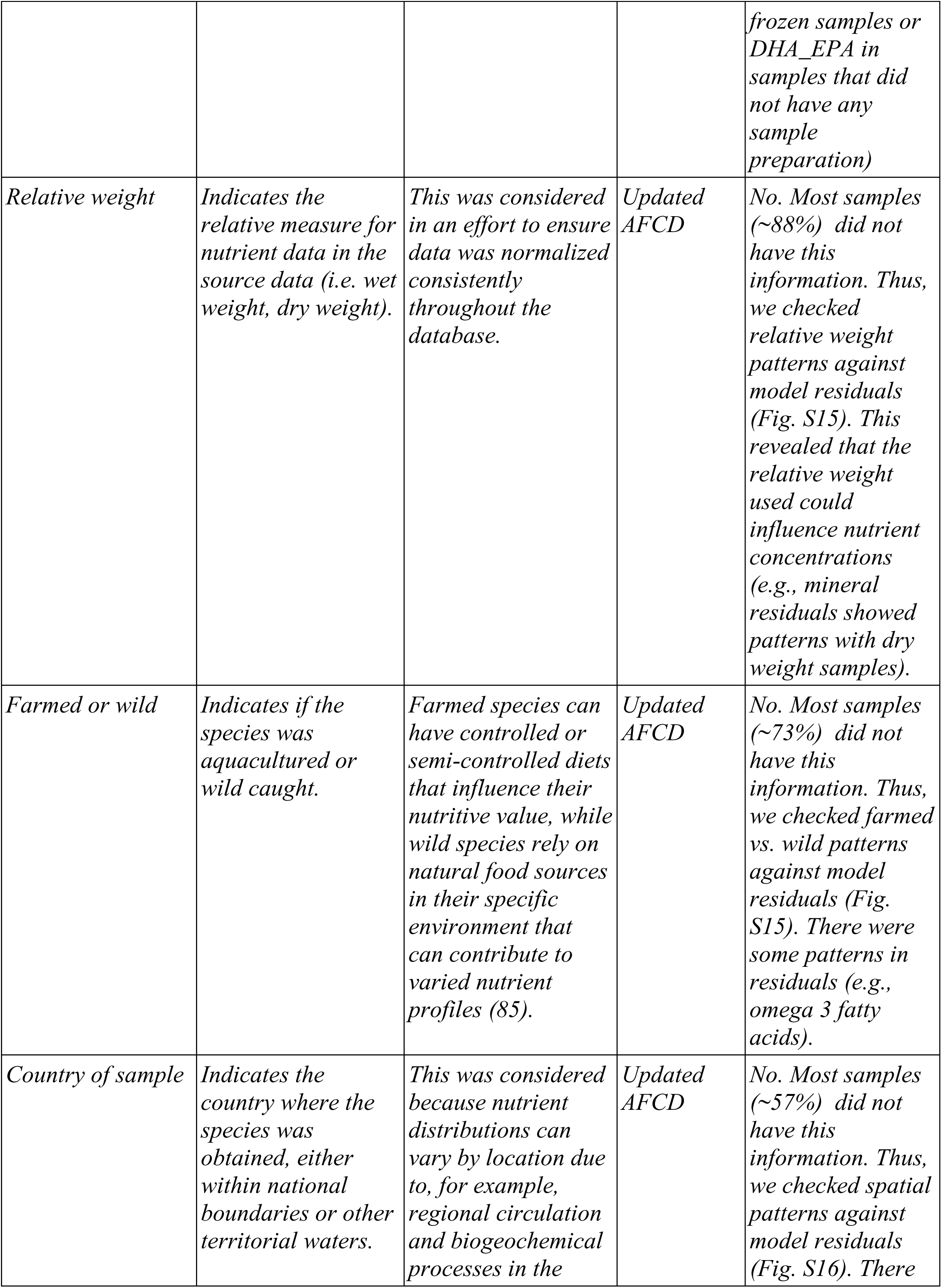
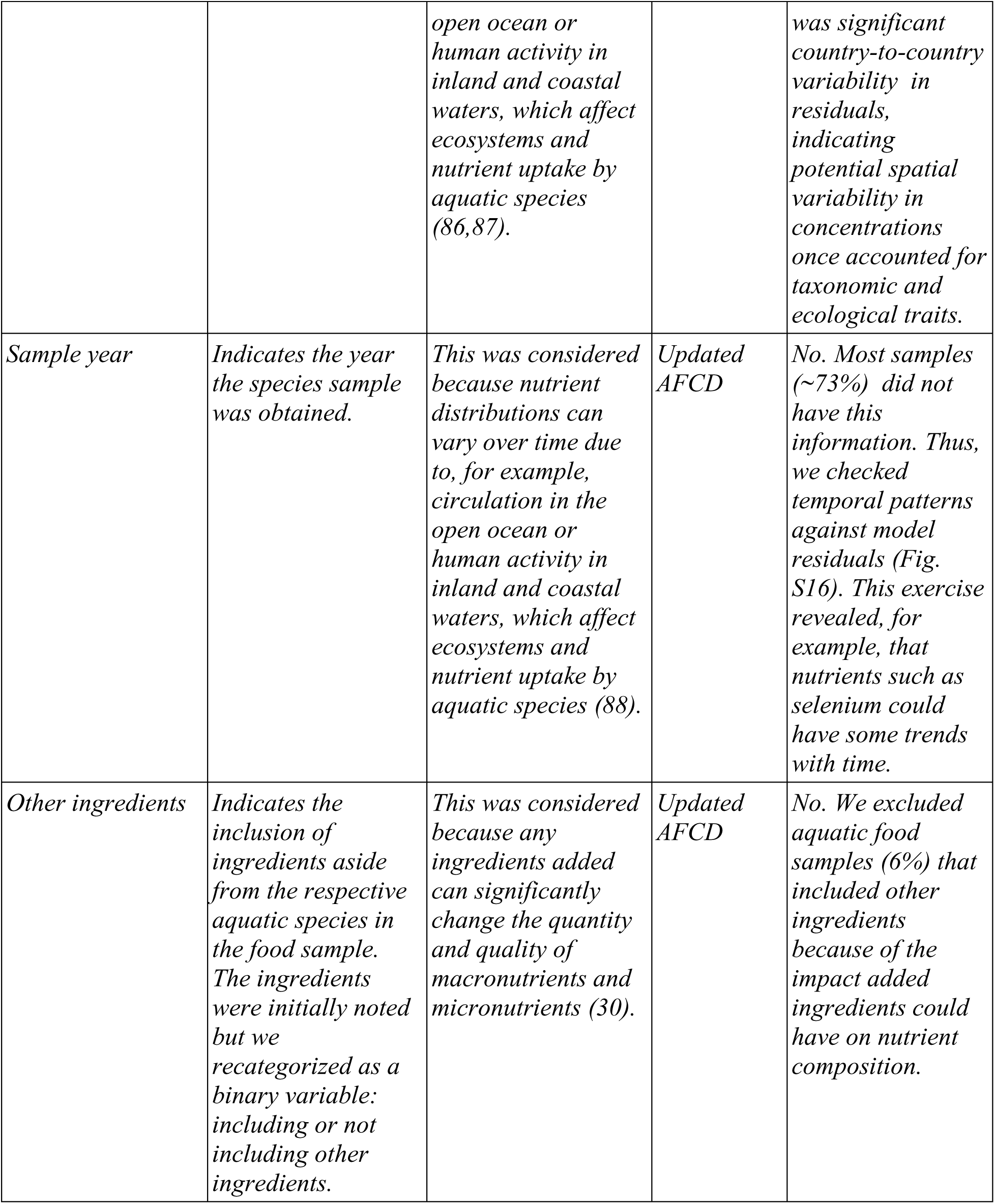
Description and source of covariates considered for our model.

**Table S2.** Nutrients and sample sizes in species-specific invertebrate data used to build invertebrate nutrient composition models. Nutrients are color-coded by whether they are minerals (red), vitamins (grey), protein (blue) or fatty acids (yellow).

| *Nutrient (units)* | *n samples in updated AFCD (without other ingredients)* | *n unique species in updated AFCD (without other ingredients)* | *n samples with all data for models* | *n sp with all data for models* | *Sufficient data to estimate parameters (based on posterior contraction values)* |
| --- | --- | --- | --- | --- | --- |
| *Iron (mg/100g)* | *966* | *331* | *706* | *226* | *Yes* |
| *Calcium (mg/100g)* | *936* | *322* | *672* | *216* | *Yes* |
| *Zinc (mg/100g)* | *848* | *278* | *663* | *198* | *Yes* |
| *Potassium (mg/100g)* | *779* | *257* | *585* | *182* | *Yes* |
| *Phosphorus (mg/100g)* | *775* | *276* | *549* | *180* | *Yes* |
| *Sodium (mg/100g)* | *757* | *243* | *572* | *174* | *Yes* |
| *Copper (mg/100g)* | *754* | *247* | *610* | *179* | *Yes* |
| *Magnesium (mg/100g)* | *658* | *216* | *508* | *155* | *Yes* |
| *Vitamin B2 (Riboflavin; mg/100g)* | *632* | *206* | *469* | *142* | *Yes* |
| *Monounsaturated fatty acids (g/100g)* | *610* | *159* | *419* | *120* | *Yes* |
| *Vitamin A (mcg/100g)* | *559* | *189* | *412* | *133* | *Yes* |
| *Vitamin B1 (Thiamin; mg/100g )* | *527* | *176* | *390 (389)* | *122 (121)* | *Yes, deleting one sample corresponding to a reproductive tissue body part that* *caused such parameter have low posterior contraction values* |
| *Vitamin C (mg/100g)* | *510* | *180* | *401 (391)* | *127 (127)* | *Yes, deleting 10 dried samples because the dried parameter had low posterior contraction* |
| *Selenium (mcg/100g)* | *481* | *165* | *417* | *131* | *Yes* |
| *DHA_EPA (g/100g)* | *457* | *133* | *320 (316)* | *101 (100)* | *Yes, deleting the only freshwater species that caused such parameter have low posterior contraction due to structure in the data* |
| *Vitamin B<sub>12</sub> (mcg/100g)* | *436* | *127* | *333* | *87* | *Yes* |
| *Protein (g/100g)* | *427* | *161* | *218* | *98* | *Yes* |
| *Manganese (mg/100g)* | *421* | *200* | *307* | *143* | *Yes* |
| *ALA (g/100g)* | *359* | *113* | *234* | *87* | *Yes, deleting five samples corresponding to dried processing and cold thermal regime which made such parameters have low posterior contraction values* |
| *Folate (mcg/100g)* | *325* | *111* | *273* | *82* | *Yes* |
| *Vitamin B5 (Pantothenic acid; mg/100g)* | *320* | *101* | *259* | *46* | *Yes* |
| *Chromium*<br>*(mcg/100g)* | 298 | 141 | 246<br>(245) | 108<br>(107) | *Yes, deleting one sample for a demersal species. The demersal parameter was informed by the prior (low posterior contraction) due to only one sample in the demersal category. After deleting that sample, the model had good posterior contraction values* |
| *Vitamin B<sub>6</sub>*<br>*(mg/100g)* | 280 | 93 | 244 | 72 | *Yes* |
| *Iodine*<br>*(mcg/100g)* | 251 | 74 | 234 | 62 | *Yes* |
| *Vitamin B<sub>3</sub>*<br>*(Niacin;*<br>*mg/100g)* | 174 | 41 | 167 | 38 | *Yes but using slightly stronger (i.e., student_t (3,0,1)) priors for the standard deviation component. The full model did not include effect sizes for body parts because there were only muscle tissue samples* |
| *Omega 3 fatty acids (g/100g)* | 119 | 74 | 90 | 53 | *Yes, but the model without covariates was preferred (probably due to limited sample size)* |
| *Omega 6 fatty acids (g/100g)* | 103 | 66 | 75 | 48 | *Yes, but the model without covariates was preferred (probably due to limited sample size)* |
| *Vitamin D*<br>*(mcg/100g)* | 29 | 18 | 19 | 12 | *No, low posterior contraction and* *convergence issues*<br>*due to limited sample*<br>*size* |
| *Vitamin E*<br>*(mg/100g)* | *28* | *22* | *19* | *16* | *No, due to limited*<br>*sample size* |

**Table S3.** Recommended daily nutritional intakes (RNI) used for each nutrient (i.e., average RNI across demographic groups).

| **Nutrient** | **RNI** |
| --- | --- |
| *Iron* | *11.97 mg/d* |
| *Calcium* | *1025.45 mg/d* |
| *Zinc* | *8.91 mg/d* |
| *Potassium* | *2552.73 mg/d* |
| *Phosphorous* | *783.41 mg/d* |
| *Sodium* | *1303.64 mg/d* |
| *Copper* | *840.00 mcg/d* |
| *Magnesium* | *299.77 mg/d* |
| *Selenium* | *50.23 mcg/d* |
| *Manganese* | *1.85 mg/d* |
| *Chromium* | *26.35 mcg/d* |
| *Iodine* | *167.73 mcg/d* |
| *MUFA* | *48.89 g/d* |
| *DHA_EPA* | *433.33 mg/d* |
| *ALA* | *1.20 g/d* |
| *Omega 3 fatty acids* | *1633.33 mg/d* |
| *Omega 6 fatty acids* | *11.86 g/d* |
| *Protein* | *47.82 g/d* |
| *Vitamin A* | *767.73 mcg/day* |
| *Vitamin C* | *73.86 mg/d* |
| *Vitamin B12* | *2.13 mcg/d* |
| *Vitamin B6* | *1.33 mg/d* |
| *Vitamin B1* | *1.05 mg/d* |
| *Vitamin B9* | *381.59 mcg/d* |
| *Vitamin B2* | *1.11 mg/d* |
| *Vitamin B3* | *13.59 mg/d* |
| *Vitamin B5* | *4.80 mg/d* |
| *Vitamin D* | *15.00 mcg/d* |

**Table S4.** Model comparison through leave-out-one cross-validation. Values for each nutrient are the expected log predictive density differences (and SE) where 0 indicates that the model was preferred in terms of predictive accuracy. The last row shows the sum (across nutrients) of the leave-out-one information criterias for each model (null, only taxonomic hierarchy or full model). This shows the full model was substantially preferred (lower LOOIC).

| **Nutrient** | **Null model** | **Only taxonomic hierarchy** | **Full model** |
| --- | --- | --- | --- |
| *Iron* | -117.7 (22.3) | -13.9 (10.8) | 0 |
| *Calcium* | -201.1 (25.7) | -33.5 (12.9) | 0 |
| *Zinc* | -86.5 (23.3) | -0.2 (7.2) | 0 |
| *Potassium* | -125.1 (28.8) | -29.7 (21.5) | 0 |
| *Phosphorous* | -185.8 (60.7) | -36.7(14.6) | 0 |
| *Sodium* | -197.7 (28.3) | -51.9 (17.1) | 0 |
| *Copper* | -149.0 (26.1) | -18.5 (11.5) | 0 |
| *Magnesium* | -179.5 (30.1) | -44.4 (27.9) | 0 |
| *Selenium* | -115.6 (23.4) | -17.8 (13.9) | 0 |
| *Manganese* | -44.2 (14.2) | -3.3 (8.8) | 0 |
| *Chromium* | -56.1 (15.1) | -10.2 (7.5) | 0 |
| *Iodine* | -178.9 (16.1) | -14.5 (7.3) | 0 |
| *MUFA* | -185.2 (28.7) | -79.0 (25.5) | 0 |
| *DHA_EPA* | -27.4 (13.3)/ -25.3 (13.9) | -11.4 (7.8)/ -18.5 (10.6) | 0 |
| *ALA* | -51.3 (31.1) | 0 | -2.0 (10.3) |
| *Omega 3 fatty acids* | -1.5 (5.8) | 0 | -3.5 (6.5) |
| *Omega 6 fatty acids* | -35.8 (10.1) | 0 | -1.6 (5.4) |
| *Protein* | -57.6 (15.3) | -9.9 (10.8) | 0 |
| *Vitamin A* | -84.8 (13.6) | -6.0 (7.9) | 0 |
| *Vitamin C* | -120.1 (17.0) | -4.4 (7.0) | 0 |
| *Vitamin B12* | -239.9 (22.8) | 0 | -7.6 (6.2) |
| *Vitamin B6* | -78.4 (15.6) | -5.6 (6.9) | 0 |
| *Vitamin B1* | -74.0 (14.6) | -4.2 (9.0) | 0 |
| *Vitamin B9* | -159.0 (18.6) | 0 | -5.2 (6.4) |
| *Vitamin B2* | -153.1(18.1) | -27.8 (10.6) | 0 |
| *Vitamin B3* | -164.2 (12.8) | -73.7 (7.2) | 0 |
| *Vitamin B5* | -92.2 (18.8) | -24.7 (10.6) | 0 |
| **Summed LOOIC** | 65113.07 | 59711.52 | 58704.13 |

## Supplementary Text

Grateful citation report: We used R version 4.2.1 (R Core Team 2022a) and the following R packages: arrow v. 17.0.0.1 (Richardson et al. 2024), bayesplot v. 1.11.1 (Gabry et al. 2019; Gabry and Mahr 2024), brms v. 2.18.0 (Bürkner 2017, 2018, 2021), broom.mixed v. 0.2.9.4 (Bolker and Robinson 2022), coda v. 0.19.4.1 (Plummer et al. 2006), data.table v. 1.16.2 (Barrett et al. 2024), dggridR v. 3.1.0 (Barnes and Sahr 2024), DHARMa v. 0.4.7 (Hartig 2024), GGally v. 2.2.1 (Schloerke et al. 2024), ggpubr v. 0.6.0 (Kassambara 2023), ggrepel v. 0.9.2 (Slowikowski 2022), ggridges v. 0.5.6 (Wilke 2024), grid v. 4.2.1 (R Core Team 2022b), gsl v. 2.1.8 (Hankin 2006), lme4 v. 1.1.35.5 (Bates et al. 2015), matrixStats v. 1.4.1 (Bengtsson 2024), parallel v. 4.2.1 (R Core Team 2022c), plyr v. 1.8.9 (Wickham 2011), reshape2 v. 1.4.4 (Wickham 2007), rnaturalearth v. 1.0.1 (Massicotte and South 2023), rnaturalearthdata v. 1.0.0 (South, Michael, and Massicotte 2024), rstan v. 2.32.6 (Stan Development Team 2024), sf v. 1.0.19 (Pebesma 2018; Pebesma and Bivand 2023), tidyverse v. 2.0.0 (Wickham et al. 2019), VIM v. 6.2.2 (Kowarik and Templ 2016), viridis v. 0.6.5 (Garnier et al. 2024).

## References and Notes

1. Passarelli, S., Free, C. M., Shepon, A., Beal, T., Batis, C., & Golden, C. D. (2024). Global estimation of dietary micronutrient inadequacies: a modelling analysis. The Lancet Global Health, 12(10), e1590–e1599.

2. Horton, S., Alderman, H., & Rivera, J. A. (2008). Hunger and malnutrition. Copenhagen: Copenhagen Consensus Center.

3. Smith, M. R., & Myers, S. S. (2018). Impact of anthropogenic CO2 emissions on global human nutrition. Nature Climate Change, 8(9), 834–839.

4. Cheung, W. W., Maire, E., Oyinlola, M. A., Robinson, J. P., Graham, N. A., Lam, V. W., … & Hicks, C. C. (2023). Climate change exacerbates nutrient disparities from seafood. Nature Climate Change, 1–8.

5. Bar-On, Y. M., Phillips, R., & Milo, R. (2018). The biomass distribution on Earth. Proceedings of the National Academy of Sciences, 115(25), 6506–6511.

6. Appeltans, W., Ahyong, S. T., Anderson, G., Angel, M. V., Artois, T., Bailly, N., … & Costello, M. J. (2012). The magnitude of global marine species diversity. Current biology, 22(23), 2189–2202.

7. Harper, S., Adshade, M., Lam, V. W., Pauly, D., & Sumaila, U. R. (2020). Valuing invisible catches: Estimating the global contribution by women to small-scale marine capture fisheries production. PloS one, 15(3), e0228912.

8. Paine, R. T. (1969). A note on trophic complexity and community stability. The American Naturalist. 103 (929): 91–93. http://www.jstor.org/stable/2459472

9. Melnychuk, M. C., Clavelle, T., Owashi, B., & Strauss, K. (2017). Reconstruction of global ex-vessel prices of fished species. ICES Journal of Marine Science, 74(1), 121–133.

10. Golden, C. D., Koehn, J. Z., Shepon, A., Passarelli, S., Free, C. M., Viana, D. F., … & Thilsted, S. H. (2021). Aquatic foods to nourish nations. Nature, 598(7880), 315–320.

11. Zamborain-Mason, J., Viana, D., Nicholas, K., Jackson, E. D., Koehn, J. Z., Passarelli, S., … & Golden, C. D. (2023). A Decision Framework for Selecting Critically Important Nutrients from Aquatic Foods. Current Environmental Health Reports, 1–12.

12. Bernhardt, J. R., & O’Connor, M. I. (2021). Aquatic biodiversity enhances multiple nutritional benefits to humans. Proceedings of the National Academy of Sciences, 118(15), e1917487118.

13. Costello, C., Cao, L., Gelcich, S., Cisneros-Mata, M. Á., Free, C. M., Froehlich, H. E., … & Lubchenco, J. (2020). The future of food from the sea. Nature, 588(7836), 95–100.

14. Anderson, S. C., Mills Flemming, J., Watson, R., & Lotze, H. K. (2011). Rapid global expansion of invertebrate fisheries: trends, drivers, and ecosystem effects. PLOS one, 6(3), e14735.

15. Blasco, G. D., Ferraro, D. M., Cottrell, R. S., Halpern, B. S., & Froehlich, H. E. (2020). Substantial gaps in the current fisheries data landscape. Frontiers in Marine Science, 7, 612831.

16. Collier, K. J., Probert, P. K., & Jeffries, M. (2016). Conservation of aquatic invertebrates: concerns, challenges and conundrums. Aquatic Conservation: Marine and Freshwater Ecosystems, 26(5), 817–837.

17. Eisenhauer, N., Bonn, A., & A. Guerra, C. (2019). Recognizing the quiet extinction of invertebrates. Nature communications, 10(1), 50.

18. Chen, E. Y. S. (2021). Often overlooked: Understanding and meeting the current challenges of marine invertebrate conservation. Frontiers in Marine Science, 8.

19. Vaitla, B., Collar, D., Smith, M. R., Myers, S. S., Rice, B. L., & Golden, C. D. (2018). Predicting nutrient content of ray-finned fishes using phylogenetic information. Nature Communications, 9(1), 3742.

20. Hicks, C. C., Cohen, P. J., Graham, N. A., Nash, K. L., Allison, E. H., D’Lima, C., … & MacNeil, M. A. (2019). Harnessing global fisheries to tackle micronutrient deficiencies. Nature, 574(7776), 95–98.

21. Heilpern, S. A., DeFries, R., Fiorella, K., Flecker, A., Sethi, S. A., Uriarte, M., & Naeem, S. (2021). Declining diversity of wild-caught species puts dietary nutrient supplies at risk. Science Advances, 7(22), eabf9967

22. Robinson, J. P., Nash, K. L., Blanchard, J. L., Jacobsen, N. S., Maire, E., Graham, N. A., … & Hicks, C. C. (2022). Managing fisheries for maximum nutrient yield. Fish and Fisheries, 23(4), 800–811.

23. Gentry, R.R., Froehlich, H.E., Grimm, D. et al. Mapping the global potential for marine aquaculture. Nat Ecol Evol 1, 1317–1324 (2017). 10.1038/s41559-017-0257-9

24. Gephart, J.A., Henriksson, P.J.G., Parker, R.W.R. et al. Environmental performance of blue foods. Nature 597, 360–365 (2021). 10.1038/s41586-021-03889-2

25. Pauly D., Zeller D., Palomares M.L.D. (Editors), 2020. Sea Around Us Concepts, Design and Data (seaaroundus.org)

26. FAO, “Fishery and Aquaculture Statistics. Global production by production source 1950-2020 (FishStatJ).” (FishStatJ, v2022.1.0, FAO Fisheries and Aquaculture Division, Rome, Italy, 2022); www.fao.org/fishery/statistics/software/fishstatj/en

27. Palomares, M. L. D., Pauly. D. (2023). SeaLifeBase. World Wide Web electronic publication (sealifebase.org)

28. Lall, S. P. (2022). The minerals. In Fish nutrition (pp. 469–554). Academic Press.

29. Jobling, M., & Bendiksen, E. Å. (2003). Dietary lipids and temperature interact to influence tissue fatty acid compositions of Atlantic salmon, Salmo salar L., parr. Aquaculture Research, 34(15), 1423–1441.

30. Sampels, S. (2015). The effects of processing technologies and preparation on the final quality of fish products. Trends in Food Science & Technology, 44(2), 131–146. 10.1016/j.tifs.2015.04.003

31. Byrd, K. A., Thilsted, S. H., & Fiorella, K. J. (2021). Fish nutrient composition: a review of global data from poorly assessed inland and marine species. Public Health Nutrition, 24(3), 476–486. 10.1017/S1368980020003857

32. Roos, N., Islam, M. M., and Thilsted, S. H. (2003). Small indigenous fish species in Bangladesh: Contribution to vitamin A, calcium and iron intakes. The Journal of Nutrition, 133(11), 4021S–4026S. 10.1093/jn/133.11.4021S

33. Fitriana, R. (2021) Study on the gender analysis of blue swimming crab fishery in Lampung. Environmental Defense Fund.

34. OBIS (2023) Ocean Biodiversity Information System. Intergovernmental Oceanographic Commission of UNESCO. https://obis.org.

35. Turner, A. D., Lewis, A. M., Bradley, K., & Maskrey, B. H. (2021). Marine invertebrate interactions with harmful algal blooms–implications for one health. Journal of invertebrate pathology, 186, 107555.

36. Elsler, L. G., Zamborain-Mason, J., & Golden, C. D. (2024). Seven strategies advancing climate-smart aquatic food systems to improve nutritional resilience. One Earth, 7(10), 1665–1669.

37. Bogard, J. R., Thilsted, S. H., Marks, G. C., Wahab, M. A., Hossain, M. A., Jakobsen, J., & Stangoulis, J. (2015). Nutrient composition of important fish species in Bangladesh and potential contribution to recommended nutrient intakes. Journal of Food Composition and Analysis, 42, 120–133

38. Imai, K., & Ratkovic, M. (2014). Covariate balancing propensity score. Journal of the Royal Statistical Society Series B: Statistical Methodology, 76(1), 243–263.

39. Liu, X., Huang, L., Lim, L., Fazhan, H., & Tan, K. (2024). The impact of elevated temperature on the macro-nutrients of commercially important marine bivalves: the implication of ocean warming. Critical Reviews in Food Science and Nutrition, 1–8.

40. Robinson, J. P., Maire, E., Bodin, N., Hempson, T. N., Graham, N. A., Wilson, S. K., … & Hicks, C. C. (2022). Climate-induced increases in micronutrient availability for coral reef fisheries. One Earth, 5(1), 98–108.

41. Seto, K. L., Friedman, W. R., Eurich, J. G., Gephart, J. A., Zamborain-Mason, J., Sharp, M., … & Golden, C. D. (2024). Characterizing pathways of seafood access in small island developing states. Proceedings of the National Academy of Sciences, 121(7), e2305424121.

42. Nguyen, B. N., Shen, E. W., Seemann, J., Correa, A. M., O’Donnell, J. L., Altieri, A. H., … & Leray, M. (2020). Environmental DNA survey captures patterns of fish and invertebrate diversity across a tropical seascape. Scientific reports, 10(1), 6729.

43. Lee Son, G. S., Romain, S., Rose, C. S., Moore, B. J., Magrane, K. A., Packer, P. S., & Wallace, F. R. (2023). Development of electronic monitoring (EM) computer vision systems and machine learning algorithms for automated catch accounting in Alaska fisheries.

44. Kolbusz, J., Langlois, T., Pattiaratchi, C., & de Lestang, S. (2021). Using an oceanographic model to investigate the mystery of the missing puerulus. Biogeosciences Discussions, 2021, 1–37.

45. Kleiber, D., Harris, L.M. and Vincent, A.C.J. (2015), Gender and small-scale fisheries: a case for counting women and beyond. Fish Fish, 16: 547–562. 10.1111/faf.12075

46. Yoklavich, M. M., Reynolds, J., & Rosen, D. (2015). A comparative assessment of underwater visual survey tools: results of a workshop and user questionnaire.

47. Steele, R. W., MacNeil, M. A., & Hankewich, S. (2022). Direct assessment of giant red sea cucumber (Apostichopus californicus) sustainability through experimental fisheries. Canadian Journal of Fisheries and Aquatic Sciences, 80(2), 408–419.

48. Johnson, J. E., Hooper, E., & Welch, D. J. (2020). Community Marine Monitoring Toolkit: A tool developed in the Pacific to inform community-based marine resource management. Marine Pollution Bulletin, 159, 111498.

49. Tai, T. C., Sumaila, U. R., & Cheung, W. W. (2021). Ocean acidification amplifies multi-stressor impacts on global marine invertebrate fisheries. Frontiers in Marine Science, 8, 596644.

50. Eurich, J. G., Friedman, W. R., Kleisner, K. M., Zhao, L. Z., Free, C. M., Fletcher, M., … & Mills, K. E. (2024). Diverse pathways for climate resilience in marine fishery systems. Fish and Fisheries, 25(1), 38–59.

51. Tan, K., Zhang, H., Li, S., Ma, H., & Zheng, H. (2022). Lipid nutritional quality of marine and freshwater bivalves and their aquaculture potential. Critical Reviews in Food Science and Nutrition, 62(25), 6990–7014.

52. Ridlon, A.D., Wasson, K., Waters, T., Adams, J., Donatuto, J., Fleener, G., Froehlich, H., Govender, R., Kornbluth, A., Lorda, J., Peabody, B., Iv, G.P., Rumrill, S.S., Tobin, E., Zabin, C.J., Zacherl, D., Grosholz, E.D., 2021. Conservation aquaculture as a tool for imperiled marine species: Evaluation of opportunities and risks for Olympia oysters, Ostrea lurida. PLOS ONE 16, e0252810. 10.1371/journal.pone.0252810

53. Dong, Z., Liu, D., Keesing, J.K., 2010. Jellyfish blooms in China: Dominant species, causes and consequences. Marine Pollution Bulletin 60, 954–963. 10.1016/j.marpolbul.2010.04.022

54. Torri, L., Tuccillo, F., Alejandro Puente-Tapia, F., Carrara Morandini, A., Segovia, J., Nevarez-López, C.A., Leoni, V., Failla-Siquier, G., Canepa-Oneto, A., Quiñones, J., Cedeño-Posso, C., Laaz, E., Preciado, M., Schiariti, A., 2024. Jellyfish as sustainable food source: A cross-cultural study among Latin American countries. Food Quality and Preference 117, 105166. 10.1016/j.foodqual.2024.105166

55. Simard, N.S., Militz, T.A., Kinch, J. and Southgate, P.C., 2019. Artisanal, shell-based handicraft in Papua New Guinea: Challenges and opportunities for livelihoods development. Ambio, 48, pp.374–384.

56. Grantham, R., Lau, J. & Kleiber, D. Gleaning: beyond the subsistence narrative. Maritime Studies 19, 509–524 (2020). 10.1007/s40152-020-00200-3

57. Campbell, B., Hanich, Q. (2014). Fish for the future: Fisheries development and food security for Kiribati in an era of global climate change. WorldFish, Penang, Malaysia. Project Report: 2014–47.

58. Elsler, L.G., Quintana, A., Giron-Nava, A., Oostdijk, M., Stefanski, S., Guillermo, X.B., Nenadovic, M., Romero, M.J.E., Weaver, A.H., Dyck, S.R.V., Tekwa, E.W., 2022. Strong collective action enables valuable and sustainable fisheries for cooperatives. Environ. Res. Lett. 17, 105003. 10.1088/1748-9326/ac9423

59. Gephart, J. A., Agrawal Bejarano, R., Gorospe, K., Godwin, A., Golden, C. D., Naylor, R. L., … & Troell, M. (2024). Globalization of wild capture and farmed aquatic foods. Nature Communications, 15(1), 8026.

60. Food and Nutrition Board, N. A. (2011). Dietary Reference Intakes (DRIs): Recommended Dietary Allowances and Adequate Intakes.

61. Givens, D. I., & Gibbs, R. A. (2008). Current intakes of EPA and DHA in European populations and the potential of animal-derived foods to increase them: Symposium on ‘How can the n-3 content of the diet be improved?’. Proceedings of the Nutrition Society, 67(3), 273–280.

62. Lichtenstein, A.H., Appel, L.J., Brands, M., Carnethon, M., Daniels, S., Franch, H.A., Franklin, B., Kris-Etherton, P., Harris, W.S., Howard, B., et al. (2006). Diet and lifestyle recommendations revision 2006: A scientific statement from the American Heart Association Nutrition Committee. Circulation, 114, 82–96.

63. Schwingshackl, L., & Hoffmann, G. (2012). Monounsaturated fatty acids and risk of cardiovascular disease: synopsis of the evidence available from systematic reviews and meta-analyses. Nutrients, 4(12), 1989–2007.

64. Allgeier, J. E., Wenger, S., & Layman, C. A. (2020). Taxonomic identity best explains variation in body nutrient stoichiometry in a diverse marine animal community. Scientific reports, 10(1), 13718.

65. Gelman A, & Hill J. (2007). Data Analysis Using Regression and Multilevel/Hierarchical Models. Cambridge University Press.

66. Vehtari, A., Gelman, A. & Gabry, J. (2017). Practical Bayesian model evaluation using leave-one-out cross-validation and WAIC. Stat Comput 27, 1413–1432. 10.1007/s11222-016-9696-4

67. Stan Development Team. (2024). RStan: The R Interface to Stan. https://mc-stan.org/.

68. Bürkner, P.-C. (2017). brms: An R Package for Bayesian Multilevel Models Using Stan. Journal of Statistical Software 80 (1): 1–28. 10.18637/jss.v080.i01.

69. Schad, D.J., Vasishth, S., Hohenstein, S., & Kliegl, R. (2019). How to capitalize on a priori contrasts in linear (mixed) models: A tutorial. Journal of Memory and Language, 110, 104038. 10.1016/j.jml.2019.104038

70. Food and Agriculture Organization of the United Nations (2016). FAO/INFOODS Global Food Composition Database for Fish and Shellfish Version 1.0-uFiSh1.0. Rome, Italy.

71. FAO Fisheries and Resources Monitoring System (FIRMS). 2024. Marine Resource Fact Sheets. https://firms.fao.org/firms/resource/search/en [Accessed: 14 March 2024]

72. NOAA Fisheries. 2024. Sustainable Seafood: Seafood Profiles. https://www.fisheries.noaa.gov/topic/sustainable-seafood/seafood-profiles [Accessed: 18 March 2024].

73. Li, J., Xie, X., Zhu, C. et al. Edible peanut worm (Sipunculus nudus) in the Beibu Gulf: Resource, aquaculture, ecological impact and counterplan. J. Ocean Univ. China 16, 823–830 (2017). 10.1007/s11802-017-3310-z

74. Holthuis, L.B. (1980). FAO Species Catalogue. Vol. 1 Shrimps and prawns of the world. An annotated catalogue of species of interest to fisheries. FAO Fish. Synop. 125(1):271p. Rome: FAO.

75. Tsuji, Takashi. (2023). Spotting the Burrow of Salpo (Peanut Worms) on the Tidal Flats of Mactan Island, Cebu, the Philippines. 10.13140/RG.2.2.30403.37925.

76. Lambert, G., Karney, R. C., Rhee, W. Y., & Carman, M. R. (2016). Wild and cultured edible tunicates: a review. Management of Biological invasions, 7(1), 59–66.

77. Goodyear, K. L., & McNeill, S. (1999). Bioaccumulation of heavy metals by aquatic macro-invertebrates of different feeding guilds: a review. Science of the Total Environment, 229(1-2), 1–19.

78. Anderson, N. H., & Cummins, K. W. (1979). Influences of diet on the life histories of aquatic insects. Journal of the Fisheries Board of Canada, 36(3), 335–342.

79. Brown, J. H., Gillooly, J. F., Allen, A. P., Savage, V. M. & West, G. B. Toward a metabolic theory of ecology. Ecology 85, 1771–1789 (2004).

80. Thorp, J. H. (2015). Functional relationships of freshwater invertebrates. In Thorp and Covich’s Freshwater Invertebrates (pp. 65–82). Academic Press.

81. Shalders, T. C., Champion, C., Coleman, M. A., & Benkendorff, K. (2022). The nutritional and sensory quality of seafood in a changing climate. Marine Environmental Research, 176, 105590.

82. Han, C., Xiao, Y., Guo, X., Zhang, H., Ren, J., & Yang, J. (2025). Authentication of Pacific white shrimp (Litopenaeus vannamei) reared in freshwater and seawater areas using fatty acid profiles combined with chemometrics. Food Control, 168, 110897.

83. Hui, Y.H., Cross, N., Kristinsson, H.G., Lim, M.H., Nip, W.K., Siow, L.F. and Stanfield, P.S. (2006). Biochemistry of Seafood Processing. In Food Biochemistry and Food Processing, Y.H. Hui (Ed.). 10.1002/9780470277577.ch16

84. Kabahenda, M.K., Omony, P., and Hüsken, S.M.C. (2009). Post-harvest handling of low-value fish products and threats to nutritional quality: a review of practices in the Lake Victoria region. Regional Programme Fisheries and HIV/AIDS in Africa: Investing in Sustainable Solutions. The WorldFish Center. Project Report 1975. https://hdl.handle.net/20.500.12348/1463

85. David, F. (2019). A worldwide reliable indicator to differentiate wild vs. farmed Penaeid shrimps based on 207 fatty acid profiles. Food chemistry, 292, 247–252.

86. Sigman, D. M. & Hain, M. P. (2012a) The Biological Productivity of the Ocean. Nature Education Knowledge 3(10):21.

87. Tozawa, M., Nomura, D., Yamazaki, K., Kiuchi, M., Hirano, D., Aoki, S., … & Murase, H. (2024). Oceanographic factors determining the distribution of nutrients and primary production in the subpolar Southern Ocean. Progress in Oceanography, 225, 103266.

88. Sigman, D. M. & Hain, M. P. (2012b) The Biological Productivity of the Ocean: Section 2. Nature Education Knowledge 3(10):20.

## References

• Barnes, Richard, and Kevin Sahr. 2024. dggridR: Discrete Global Grids. https://CRAN.R-project.org/package=dggridR.

• Barrett, Tyson, Matt Dowle, Arun Srinivasan, Jan Gorecki, Michael Chirico, Toby Hocking, and Benjamin Schwendinger. 2024. data.table: Extension of “data.frame”. https://CRAN.R-project.org/package=data.table.

• Bates, Douglas, Martin Mächler, Ben Bolker, and Steve Walker. 2015. “Fitting Linear Mixed-Effects Models Using lme4.” Journal of Statistical Software 67 (1): 1–48. 10.18637/jss.v067.i01.

• Bengtsson, Henrik. 2024. matrixStats: Functions That Apply to Rows and Columns of Matrices (and to Vectors). https://CRAN.R-project.org/package=matrixStats.

• Bolker, Ben, and David Robinson. 2022. broom.mixed: Tidying Methods for Mixed Models. https://CRAN.R-project.org/package=broom.mixed.

• Bürkner, Paul-Christian. 2017. “brms: An R Package for Bayesian Multilevel Models Using Stan.” Journal of Statistical Software 80 (1): 1–28. 10.18637/jss.v080.i01.

• Bürkner, Paul-Christian. 2018. “Advanced Bayesian Multilevel Modeling with the R Package brms.” The R Journal 10 (1): 395–411. 10.32614/RJ-2018-017.

• Bürkner, Paul-Christian. 2021. “Bayesian Item Response Modeling in R with brms and Stan.” Journal of Statistical Software 100 (5): 1–54. 10.18637/jss.v100.i05.

• Gabry, Jonah, and Tristan Mahr. 2024. “bayesplot: Plotting for Bayesian Models.” https://mc-stan.org/bayesplot/.

• Gabry, Jonah, Daniel Simpson, Aki Vehtari, Michael Betancourt, and Andrew Gelman. 2019. “Visualization in Bayesian Workflow.” J. R. Stat. Soc. A 182: 389–402. 10.1111/rssa.12378.

• Garnier, Simon, Ross, Noam, Rudis, Robert, Camargo, et al. 2024. viridis(Lite) - Colorblind-Friendly Color Maps for r. 10.5281/zenodo.4679423.

• Hankin, Robin K. S. 2006. “Special Functions in r: Introducing the Gsl Package.” R News 6.

• Hartig, Florian. 2024. DHARMa: Residual Diagnostics for Hierarchical (Multi-Level / Mixed) Regression Models. https://CRAN.R-project.org/package=DHARMa.

• Kassambara, Alboukadel. 2023. ggpubr: “ggplot2” Based Publication Ready Plots. https://CRAN.R-project.org/package=ggpubr.

• Kowarik, Alexander, and Matthias Templ. 2016. “Imputation with the R Package VIM.” Journal of Statistical Software 74 (7): 1–16. 10.18637/jss.v074.i07.

• Massicotte, Philippe, and Andy South. 2023. rnaturalearth: World Map Data from Natural Earth. https://CRAN.R-project.org/package=rnaturalearth.

• Pebesma, Edzer. 2018. “Simple Features for R: Standardized Support for Spatial Vector Data.” The R Journal 10 (1): 439–46. 10.32614/RJ-2018-009.

• Pebesma, Edzer, and Roger Bivand. 2023. Spatial Data Science: With applications in R. Chapman and Hall/CRC. 10.1201/9780429459016.

• Plummer, Martyn, Nicky Best, Kate Cowles, and Karen Vines. 2006. “CODA: Convergence Diagnosis and Output Analysis for MCMC.” R News 6 (1): 7–11. https://journal.r-project.org/archive/.

• R Core Team. 2022a. R: A Language and Environment for Statistical Computing. Vienna, Austria: R Foundation for Statistical Computing. https://www.R-project.org/.

• R Core Team. 2022b. R: A Language and Environment for Statistical Computing. Vienna, Austria: R Foundation for Statistical Computing. https://www.R-project.org/.

• R Core Team. 2022c. R: A Language and Environment for Statistical Computing. Vienna, Austria: R Foundation for Statistical Computing. https://www.R-project.org/.

• Richardson, Neal, Ian Cook, Nic Crane, Dewey Dunnington, Romain François, Jonathan Keane, Dragoș Moldovan-Grünfeld, Jeroen Ooms, Jacob Wujciak-Jens, and Apache Arrow. 2024. arrow: Integration to “Apache” “Arrow”. https://CRAN.R-project.org/package=arrow.

• Schloerke, Barret, Di Cook, Joseph Larmarange, Francois Briatte, Moritz Marbach, Edwin Thoen, Amos Elberg, and Jason Crowley. 2024. GGally: Extension to “ggplot2”. https://CRAN.R-project.org/package=GGally.

• Slowikowski, Kamil. 2022. ggrepel: Automatically Position Non-Overlapping Text Labels with “ggplot2”. https://CRAN.R-project.org/package=ggrepel.

• South, Andy, Schramm Michael, and Philippe Massicotte. 2024. rnaturalearthdata: World Vector Map Data from Natural Earth Used in “rnaturalearth”. https://CRAN.R-project.org/package=rnaturalearthdata.

• Stan Development Team. 2024. “RStan: The R Interface to Stan.” https://mc-stan.org/.

• Wickham, Hadley. 2007. “Reshaping Data with the reshape Package.” Journal of Statistical Software 21 (12): 1–20. http://www.jstatsoft.org/v21/i12/.

• Wickham, Hadley. 2011. “The Split-Apply-Combine Strategy for Data Analysis.” Journal of Statistical Software 40 (1): 1–29. https://www.jstatsoft.org/v40/i01/.

• Wickham, Hadley, Mara Averick, Jennifer Bryan, Winston Chang, Lucy D’Agostino McGowan, Romain François, Garrett Grolemund, et al. 2019. “Welcome to the tidyverse.” Journal of Open Source Software 4 (43): 1686. 10.21105/joss.01686.

• Wilke, Claus O. 2024. ggridges: Ridgeline Plots in “ggplot2”. https://CRAN.R-project.org/package=ggridges.

